# PolyGR and polyPR knock-in mice reveal a conserved neuroprotective extracellular matrix signature in *C9orf72* ALS/FTD neurons

**DOI:** 10.1101/2023.07.17.549331

**Authors:** Carmelo Milioto, Mireia Carcolé, Ashling Giblin, Rachel Coneys, Olivia Attrebi, Mhoriam Ahmed, Samuel S. Harris, Byung Il Lee, Mengke Yang, Raja S. Nirujogi, Daniel Biggs, Sally Salomonsson, Matteo Zanovello, Paula De Oliveira, Eszter Katona, Idoia Glaria, Alla Mikheenko, Bethany Geary, Evan Udine, Deniz Vaizoglu, Rosa Rademakers, Marka van Blitterswijk, Anny Devoy, Soyon Hong, Linda Partridge, Pietro Fratta, Dario R. Alessi, Ben Davies, Marc Aurel Busche, Linda Greensmith, Elizabeth M.C. Fisher, Adrian M. Isaacs

**Affiliations:** UK Dementia Research Institute, University College London, London WC1E 6BT, UK; Department of Neurodegenerative Disease, UCL Institute of Neurology, Queen Square, London WC1N 3BG, UK; Department of Genetics, Evolution and Environment, Institute of Healthy Ageing, London, United Kingdom; Department of Neuromuscular Diseases, UCL Queen Square Institute of Neurology, London, UK; Aligning Science Across Parkinson’s (ASAP) Collaborative Research Network, Chevy Chase, MD, 20815, USA; Medical Research Council (MRC) Protein Phosphorylation and Ubiquitylation Unit, School of Life Sciences, University of Dundee, Dow Street, Dundee, DD1 5EH, UK; Wellcome Centre for Human Genetics, University of Oxford, Oxford OX3 7BN; Research Support Service, Institute of Agrobiotechnology, CSIC-Government of Navarra, Mutilva, Spain; Department of Neuroscience, Mayo Clinic, 4500 San Pablo Road S, Jacksonville, FL, 32224, USA; VIB Center for Molecular Neurology, University of Antwerp, Universiteitsplein 1, Wilrijk, 2610, Antwerp, Belgium; Department of Biomedical Sciences, University of Antwerp, Antwerp, Belgium; Maurice Wohl Clinical Neuroscience Institute, King’s College London, London SE5 9RT, UK; UCL Queen Square Motor Neuron Disease Centre, UCL Queen Square Institute of Neurology, University College London, Queen Square, London WC1N 3BG, UK; Francis Crick Institute, 1 Midland Rd, London NW1 1AT

**Keywords:** C9orf72, extracellular matrix, frontotemporal dementia, amyotrophic lateral sclerosis, collagen VI, COL6A1, TGF-β, neuronal hyperactivity

## Abstract

A GGGGCC repeat expansion in *C9orf72* is the most common genetic cause of ALS and FTD (C9ALS/FTD). The presence of dipeptide repeat (DPR) proteins, generated by translation of the expanded repeat, is a major pathogenic feature of C9ALS/FTD pathology, but their most relevant effects in a physiological context are not known. Here, we generated *C9orf72* DPR knock-in mouse models characterised by physiological expression of 400 codon-optimised polyGR or polyPR repeats, and heterozygous *C9orf72* reduction. (GR)400 and (PR)400 knock-in mice exhibit cortical neuronal hyperexcitability, age-dependent spinal motor neuron loss and progressive motor dysfunction, showing that they recapitulate key features of C9FTD/ALS. Quantitative proteomics revealed an increase in extracellular matrix (ECM) proteins in (GR)400 and (PR)400 spinal cord, with the collagen COL6A1 the most increased protein. This signature of increased ECM proteins was also present in C9ALS patient iPSC-motor neurons indicating it is a conserved feature of C9ALS/FTD. TGF-β1 was one of the top predicted regulators of this ECM signature and polyGR expression in human iPSC-neurons was sufficient to induce TGF-β1 followed by COL6A1, indicating TGF-β1 is one driver of the ECM signature. Knockdown of the TGF-β1 or COL6A1 orthologue in *Drosophila* dramatically and specifically exacerbated neurodegeneration in polyGR flies, showing that TGF-β1 and COL6A1 protect against polyGR toxicity. Altogether, our physiological *C9orf72* DPR knock-in mice have revealed a neuroprotective and conserved ECM signature in C9FTD/ALS.

## Introduction

Amyotrophic lateral sclerosis (ALS), characterised by progressive muscle weakness and atrophy, and frontotemporal dementia (FTD), characterised by personality and behavioural change or language dysfunction, are adult-onset neurodegenerative diseases and part of a disease spectrum with overlapping clinical, pathological, and genetic origins ^1, 2^. A GGGGCC (G_4_C_2_) repeat expansion in the first intron of the *C9orf72* gene ^3, 4^ is the most common genetic cause of ALS and FTD ^3, 4^ accounting for approximately 38% of familial ALS (fALS), 6% of sporadic ALS (sALS), 25% of familial FTD (fFTD) and 6% of sporadic FTD (sFTD) ^5^: collectively termed C9ALS/FTD ^6^. Three mechanisms have been proposed to induce C9ALS/FTD pathology, either by loss– or gain-of-function: (i) reduced transcription of *C9orf72*, (ii) the presence of sense and antisense repeat-containing RNA foci, (iii) expression of aberrant dipeptide repeat (DPR) proteins encoded in six frames by the hexanucleotide repeat. These mechanisms are not mutually exclusive and they are all predicted to contribute to disease to some extent. However, the presence of DPRs is a major pathogenic feature of C9ALS/FTD.

The DPRs are derived from sense and antisense repeat-containing RNAs, which are translated by repeat-associated non-ATG initiated (RAN) translation, a non-canonical protein translation mechanism that does not require an ATG start codon ^7^. RAN translation occurs in every reading frame and both RNA directions, encoding five potentially toxic DPRs: polyGA, polyGR, polyGP, polyPA and polyPR. Several studies have reported that DPR proteins accumulate in neuronal cytoplasmic inclusions in post-mortem tissues from C9ALS/FTD patients ^8–13^. Moreover, we and others have shown that DPR proteins are toxic *in vivo* and *in vitro* ^14^. The arginine-rich DPR proteins, polyGR and polyPR, are generally the most toxic species among the DPRs in several systems, including *Drosophila* ^15–17^, mammalian cells ^16–22^, and mouse models ^22–28^. Therefore, it is essential to investigate DPRs within the context of C9ALS/FTD to develop novel and effective therapeutic strategies.

Thus, several mouse models have been developed to elucidate pathomechanisms associated with C9ALS/FTD pathology ^29, 30^. *C9orf72* homozygous knockout mouse models manifest severe autoimmunity and lymphatic defects, indicating a role for C9orf72 in immune cell function, but there are no striking alterations in heterozygous knockouts which more closely resemble the degree of C9orf72 depletion in patients ^31–38^. Mouse models of *C9orf72* hexanucleotide repeat expansions have also been generated, taking advantage of either AAV-mediated delivery ^39, 40^, or bacterial artificial chromosome (BAC) integration ^33, 41–43^. Although most mouse models recapitulate features of C9ALS/FTD, there is considerable phenotypic variation between them, possibly from site of integration, which may cause local mutation, copy number or other effects ^44^. Finally, to better elucidate the role of DPRs in C9ALS/FTD, both viral and transgenic mouse models have been developed over-expressing codon-optimised constructs to synthesize only one specific DPR. Studies of these mice show that *in vivo* expression of polyGR throughout the mouse brain is toxic and results in several aspects of C9ALS/FTD pathology, including age-dependent neuronal loss, presence of cytoplasmic aggregates, development of anxiety-like behaviour and social interaction defects ^22–25^. Expression of polyPR is also toxic in mice. Similar to polyGR, polyPR mice have survival, motor and cognitive defects, hyperactivity and anxiety-like behaviour, progressive brain atrophy, and neuronal loss ^26–28^. Current *in vivo* and *in vitro* studies have identified common downstream molecular pathways that are dysregulated in C9ALS/ FTD, including autophagy, nucleocytoplasmic transport, pre-messenger RNA splicing, stress granule dynamics, DNA damage repair, mitochondrial dysfunction, nuclear pore alterations and synaptic dysfunction ^14, 45–47^. However, the mechanism(s) by which the repeat expansion causes C9ALS/FTD when DPRs are expressed at physiological levels are less clear.

The collection of mouse models produced to date have identified several potential pathomechanisms underlying C9ALS/FTD. However, the most relevant effects of DPRs in the endogenous context are not known. Thus, there is an urgent need for refined models to understand the role of individual DPRs *in vivo*. Here, we generated *C9orf72* DPR knock-in mouse models characterised by physiological DPR expression and heterozygous C9orf72 reduction, to more accurately model DPR-induced dysfunction in C9ALS/FTD and reveal new insights into molecular mechanisms contributing to disease.

## Results

### Generation of *C9orf72* polyGR and polyPR knock-in mice

As polyGR and polyPR are consistently damaging across model systems, we focused on generating polyGR and polyPR knock-in mice (**Fig. 1A**). We first generated patient-length DPRs by performing recursive directional ligation ^15, 48^ to build stretches of 400 uninterrupted codon-optimised polyGR or polyPR repeats (**Fig. 1B**) flanked by epitope tags. We then used CRISPR-Cas9 to insert these repeats, or a control eGFP sequence, immediately after, and in frame with, the endogenous mouse *C9orf72* ATG start codon, in mouse embryonic stem (ES) cells (**Supplementary Fig. 1A**). We performed targeted locus amplification to identify clones with a single insertion site, and correct targeting, which maintained the integrity of both the knock-in sequences and the adjacent mouse genome (**Supplementary Fig. 1B-1D**). Validated ES cell clones were then taken forward to generate knock-in mice using standard procedures. We developed a PCR assay to amplify across the 400 codon-optimised repeats and showed that DPR length is stable across at least five generations (**Supplementary Fig. 1E**), allowing the lines to be easily maintained and shared. To further confirm correct targeting, we measured DPR levels and, as expected, (GR)400 and (PR)400 mice selectively expressed their own DPR in brain and spinal cord at 3 months of age (**Fig. 1C and 1D**). Importantly, levels of polyGR in (GR)400 mouse cortex were similar to C9FTD/ALS patient cortex (**Supplementary Fig. 2A**), confirming expression in the physiological range. (GR)400 and (PR)400 mice exhibited a significant reduction by ∼40% of *C9orf72* at mRNA and protein levels in 3-month-old brain and spinal cord (**Fig. 1E and 1F**), as predicted by our knock-in strategy, which inserts the DPR sequence into one copy of *C9orf72*, removing expression of *C9orf72* from that allele. Finally, we confirmed eGFP expression by ELISA and *C9orf72* reduction by qPCR in brain and spinal cord of 3-month-old *C9orf72* eGFP knock-in mice (**Supplementary Fig. 2B and 2C**). These results confirm that our DPR knock-in mouse lines selectively express their specific DPR at physiological levels, in combination with *C9orf72* reduction, as predicted by our targeting strategy.

**Figure 1.**
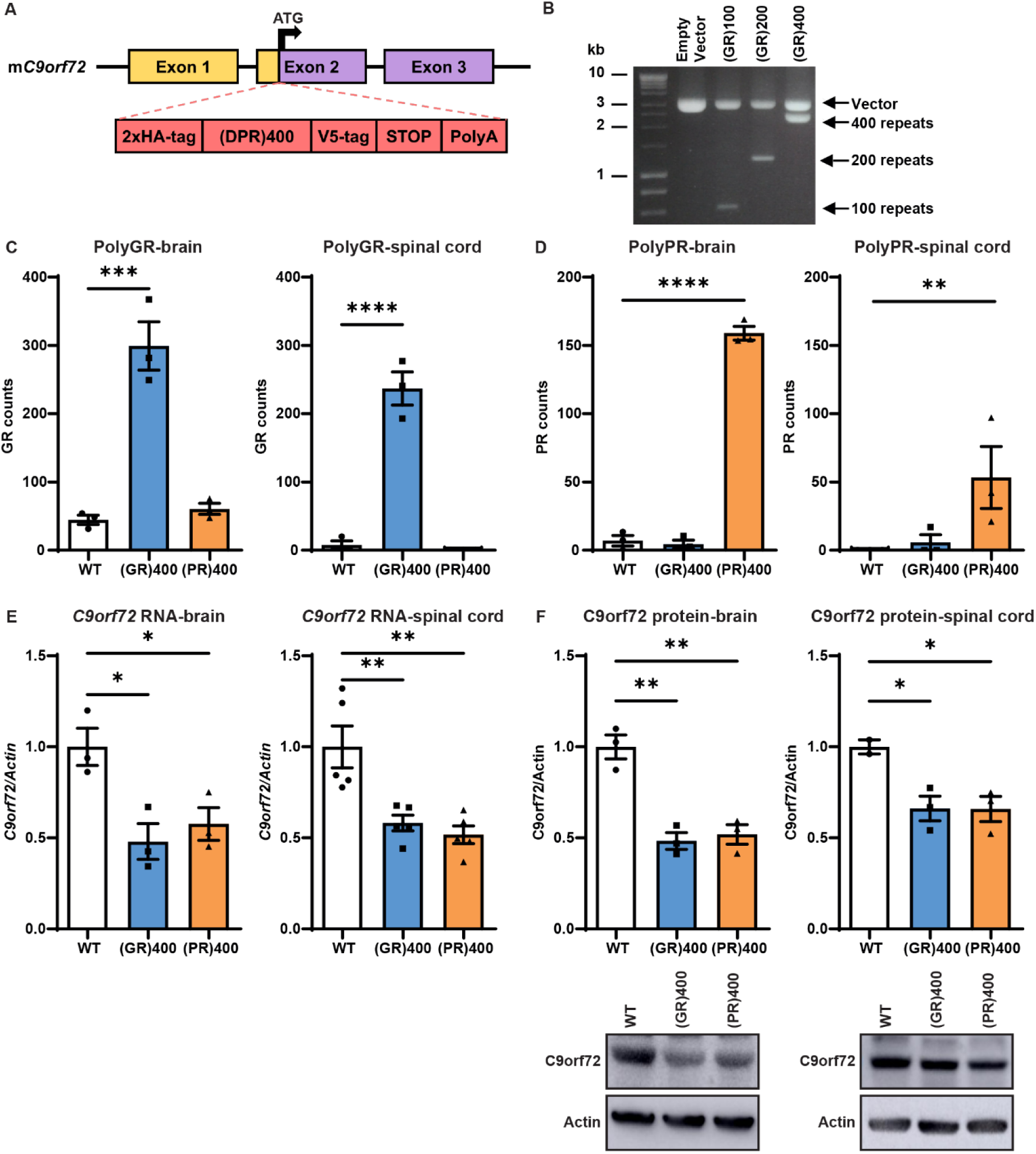
Generation of *C9orf72* polyGR and polyPR knock-in mice (A) Targeting strategy to generate (GR)400 and (PR)400 mice with the knock-in sequence inserted in exon 2 of mouse *C9orf72* immediately after, and in frame with, the endogenous ATG. Schematic shows the genomic region and the knock-in targeting construct. Exons are shown boxed, untranslated regions of exons are coloured yellow with translated regions in purple. The targeting construct (in red) contains the knock-in sequence composed of a double HA-tag, 400 codon-optimised DPRs, a V5-tag, a stop codon, and a 120 bp polyA tail. (B) Agarose gel shows generation of patient-length polyGR, using recursive directional ligation to sequentially double the repeat length up to 400 repeats. (C) Quantification of polyGR proteins in brain (left panel) and spinal cord (right panel) of WT, (GR)400, and (PR)400 mice at 3 months of age by Meso Scale Discovery (MSD) immunoassay. Graph, mean ± SEM, n = 3 mice per genotype, one-way ANOVA, Bonferroni’s multiple comparison, ***p < 0.001, ****p < 0.0001. (D) Quantification of polyPR proteins in brain (left panel) and spinal cord (right panel) of WT, (GR)400, and (PR)400 mice at 3 months of age by MSD immunoassay. Graph, mean ± SEM, n = 3 mice per genotype, one-way ANOVA, Bonferroni’s multiple comparison, **p < 0.01, ****p < 0.0001. (E) Quantitative PCR analysis of *C9orf72* transcript levels normalised to *β-actin* in brain (left panel) and spinal cord (right panel) of WT, (GR)400, and (PR)400 mice at 3 months of age. Graph, mean ± SEM, n = 3 mice per genotype, one-way ANOVA, Bonferroni’s multiple comparison, *p < 0.05, **p < 0.01, ***p < 0.001. (F) Western blotting analysis of C9orf72 protein levels in brain (left panel) and spinal cord (right panel) of WT, (GR)400, and (PR)400 mice at 3 months of age. β-actin is shown as loading control. Graph, mean ± SEM, n = 3 mice per genotype, one-way ANOVA, Bonferroni’s multiple comparison, *p < 0.05, **p < 0.01.

**Figure 2.**
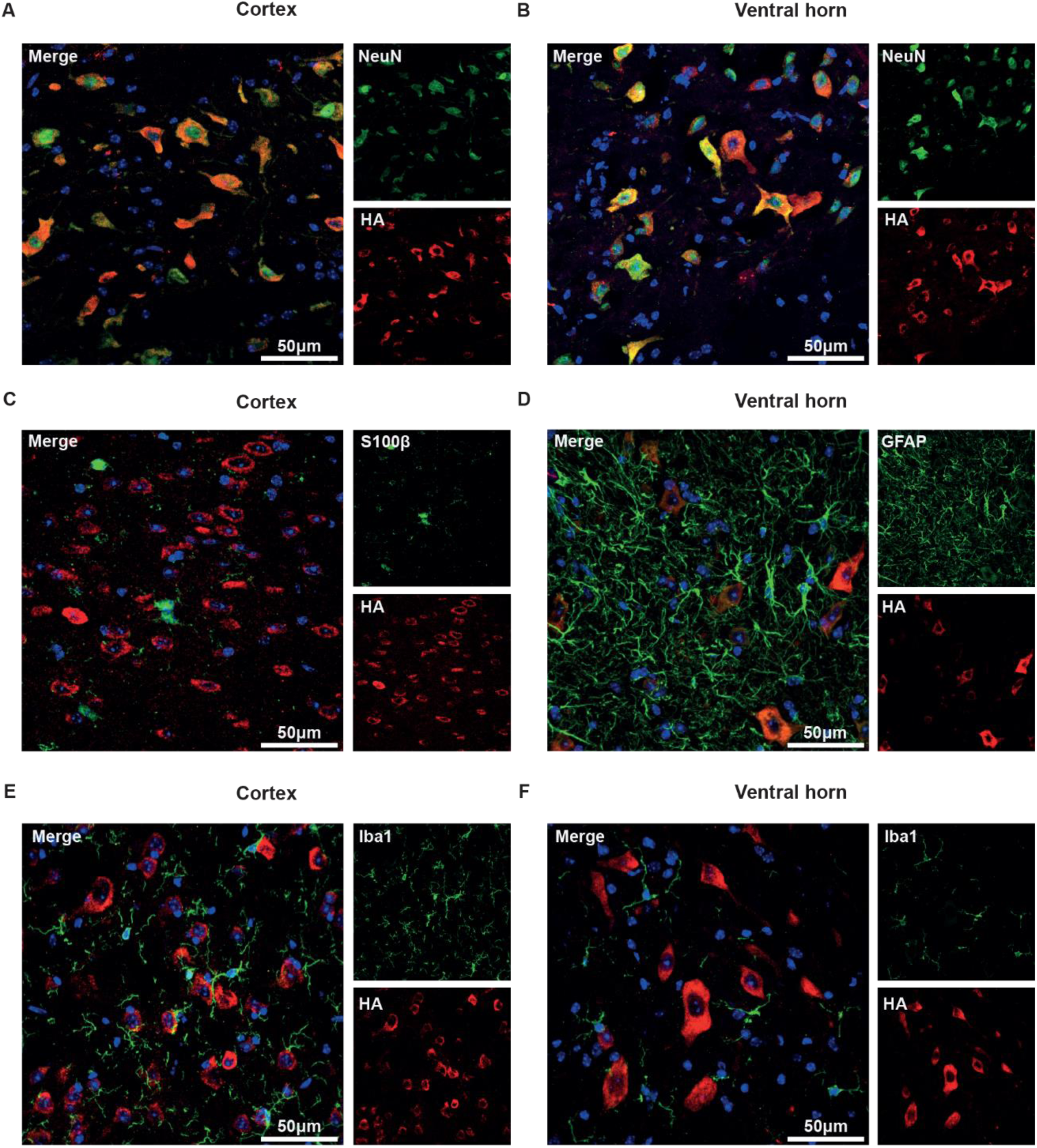
PolyGR is predominantly expressed in neurons (A-B) Representative confocal images of immunofluorescence staining showing colocalization between neuronal marker NeuN (green) and HA-tag (red) in (GR)400 (A) mouse brain cortex and (B) lumbar spinal cord ventral horn at 6 months of age. (C-D) Representative confocal images of immunofluorescence staining showing absence of colocalization between the astrocytic markers S100β (C) or GFAP (D) (green) and HA-tag (red) in (GR)400 mouse (C) brain cortex and (D) lumbar spinal cord ventral horn at 6 months of age. (E-F) Representative confocal images of immunofluorescence staining showing absence of colocalization between the microglial marker Iba1 (green) and HA-tag (red) in (GR)400 mouse (E) brain cortex and (F) lumbar spinal cord ventral horn at 6 months of age. DAPI (blue) stains nuclei.

### PolyGR is predominantly expressed in neurons

Having established DPRs are translated in brain and spinal cord, we used the HA-tag to visualise them by immunostaining. PolyGR showed a clear cytoplasmic localisation. However, we did not observe polyPR staining in the same conditions, perhaps due to inaccessibility of the epitope tag and therefore we focussed on characterising polyGR in more detail. We found that polyGR shows widespread neuronal expression, co-localising with a neuronal marker (NeuN) in cortex and in the ventral horn of lumbar spinal cord in 6-month-old mice (**Fig. 2A and 2B**). Interestingly, at the same time-point, we did not observe co-localisation of polyGR with astrocytic S100β, GFAP, or microglial markers (Iba1) in cortex and ventral horn of lumbar spinal (**Fig. 2C-F**). These results show that in our knock-in mice, polyGR is neuronally expressed in FTD/ALS relevant regions.

### (GR)400 knock-in mice exhibit cortical hyperexcitability without TDP-43 mislocalisation or neuronal loss

Next, we assessed whether polyGR or polyPR expression in the brain is associated with pathological features of ALS and FTD. We did not observe astrogliosis in the motor cortex of (GR)400 and (PR)400 mice up to 12 months of age (**Supplementary Fig. 3A**). As microglia-mediated neuroinflammation has been observed in ALS and FTD and is associated with C9orf72 function, we assessed levels of two microglial markers, Iba1 and CD68. While Iba1 stains microglia cells and processes and thus provided an index of microglial density, lysosomal antigen CD68 served as an indirect measure of microglial phagocytic activity. Our staining revealed no alterations in microglial density or the percentage of CD68-positive microglia in the motor cortex of (GR)400 and (PR)400 mice at 12 months of age (**Supplementary Fig. 3C**). We then explored TAR DNA binding protein 43 (TDP-43) pathology, a typical hallmark in the majority of ALS patients. TDP-43 was predominantly nuclear in the cortex of (GR)400 and (PR)400 mice, without mis-localisation to the cytoplasm or inclusion formation, suggesting TDP-43 was not altered at 12 months of age (**Supplementary Fig. 3E**). We also evaluated whether expression of polyGR or polyPR caused neuronal loss. First, we measured the density of NeuN-positive neurons in the whole cortex and specifically in the motor cortex, which revealed that NeuN+ neuron density was not altered in (GR)400 and (PR)400 mice at 12 months of age (**Fig. 3A**). Then, we quantified CTIP2-positive upper motor neuron (MN) density in layer V of the motor cortex, which is particularly vulnerable to cell death in ALS. Our quantification indicated there was no significant loss of CTIP2 neurons in 12-month old (GR)400 and (PR)400 mice (**Fig. 3B**). These results suggest that, up to 12 months of age, polyGR and polyPR expression in the brain is not associated with pathological features found in ALS and FTD, including astrogliosis, microgliosis, TDP-43 pathology, and neuronal loss.

**Figure 3.**
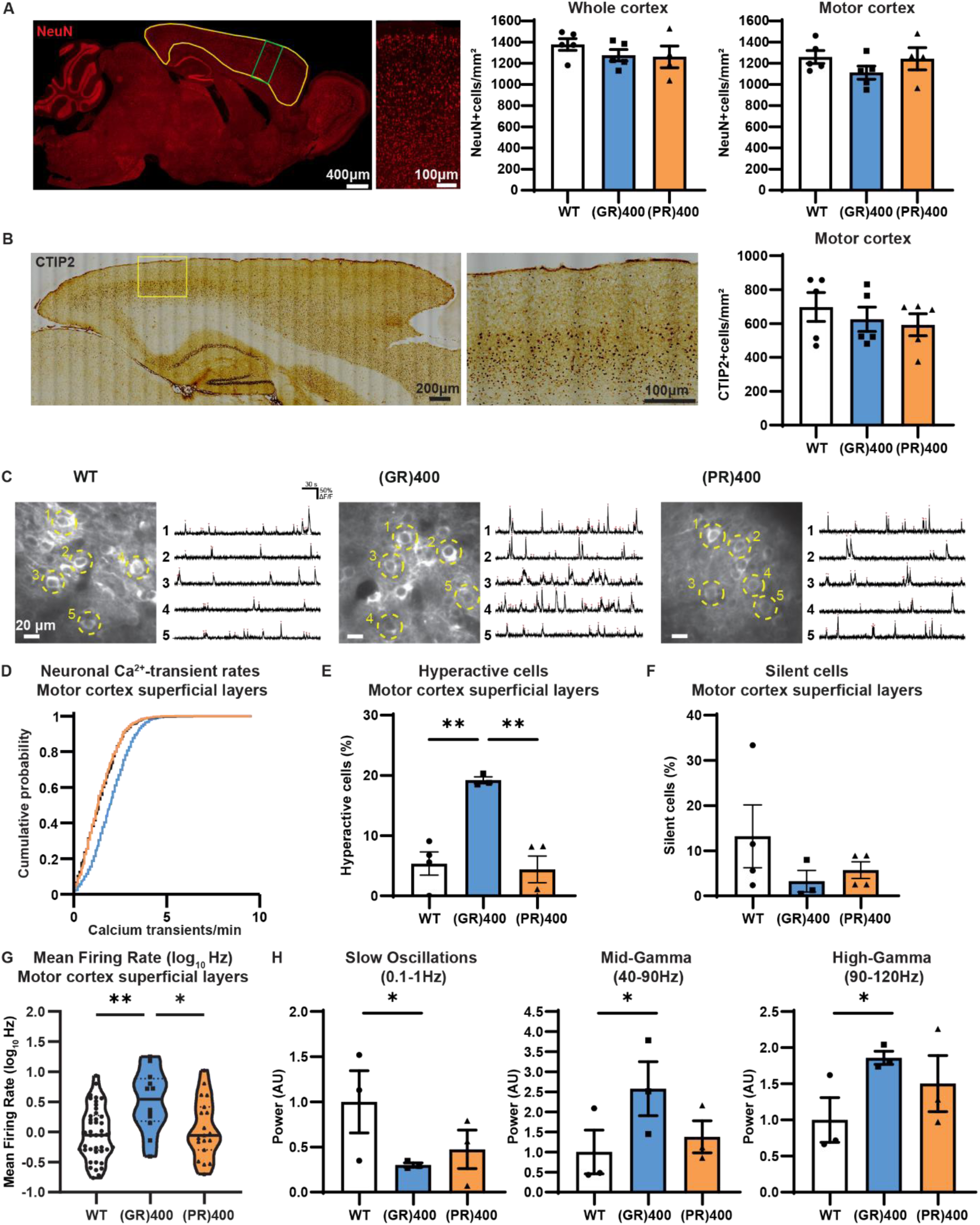
(GR)400 knock-in mice exhibit cortical hyperexcitability without neuronal loss (A) Representative image of the cortex (area delineated in yellow) and enhanced magnification of the motor cortex (area delineated in green), and quantification of NeuN-positive (red) cell density in the whole cortex and in motor cortex in WT, (GR)400, and (PR)400 mice at 12 months of age. Graph, mean ± SEM, n = 5 mice per genotype, one-way ANOVA, Bonferroni’s multiple comparison. (B) Representative image of the cortex, enhanced magnification of the motor cortex (area delineated in yellow), and quantification of CTIP2-positive upper motor neuron density in layer V of the motor cortex in WT, (GR)400, and (PR)400 mice at 12 months of age. Graph, mean ± SEM, n = 5 mice per genotype, one-way ANOVA, Bonferroni’s multiple comparison. (C) Example of *in vivo* two-photon fluorescence images of jRCamP1b-expressing superficial layer neurons in motor cortex and spontaneous Ca^2+^ activity from five example neurons in WT (left panel), GR(400) (centre panel), and PR(400) (right panel) mice at 15-19 months of age. (D) Cumulative distribution plot displaying neuronal Ca^2+^-transient rates across animals in motor cortex superficial layers of WT (732 cells, 4 mice), (GR)400 (1405 cells, 3 mice), and (PR)400 (946 cells, 4 mice) mice at 15-19 months of age. (E) Percentage of hyperactive (>3 Ca^2+^-transients per minute) neurons in motor cortex superficial layers of WT (n = 4 mice), (GR)400 (n = 3 mice), and (PR)400 (n = 4 mice) mice at 15-19 months of age. Graph, mean ± SEM, one-way ANOVA, Tukey’s multiple comparisons, **p < 0.01. (F) Percentage of silent (0 Ca^2+^-transients per min) neurons in motor cortex superficial layers of WT (n = 4 mice), (GR)400 (n = 3 mice), and (PR)400 (n = 4 mice) mice at 15-19 months of age. Graph, mean ± SEM, one-way ANOVA, Tukey’s multiple comparisons. (G) Mean firing rate (log_10_Hz) of neurons in motor cortex superficial layers of WT, (GR)400, and (PR)400 mice at 15-19 months of age; black lines indicate medians, dashed lines indicate quartiles. Graph, n = 3 mice per genotype, one-way ANOVA, Tukey’s multiple comparison, *p < 0.05, **p < 0.01. (H) Local-field potential (LFP) power in the slow-wave frequency band (left panel), mid gamma frequency band (centre panel), and high gamma frequency band (left panel) in WT, (GR)400, and (PR)400 mice at 15-19 months of age. Graph, mean ± SEM, n = 3 mice per genotype, one-way ANOVA, Fisher’s least significant difference procedure, *p < 0.05.

We therefore investigated whether functional deficits were present, which would provide insight into the early brain changes in FTD and ALS. We conducted *in vivo* two-photon calcium imaging of neurons in superficial and deep layers of the motor cortex of (GR)400 and (PR)400 mice as well as WT controls using the red-shifted genetically encoded calcium indicator jRCaMP1b (**Fig. 3C**). We found that, in (GR)400 mice, there was an increase in the fraction of abnormally hyperactive neurons in superficial layers, but not in layer 5 where spontaneous neuronal activity was comparable to (PR)400 and WT mice (**Fig. 3D-F and Supplementary Fig. 4A-C**). To further validate these layer-specific neuronal impairments, we performed high-density *in vivo* Neuropixel recordings in a subset of the same mice and in an additional cohort to assess single-unit and population neuronal activity. These experiments confirmed that motor cortex neurons in superficial layers in (GR)400 mice were hyperexcitable relative to those in (PR)400 and WT mice, (**Fig. 3G and Supplementary Fig. 4D**). Furthermore, evaluation of population local field potential (LFP) across all motor cortex laminae was suggestive of a concurrent reduction in slow wave activity power, and an increase in gamma-frequency band power, in (GR)400 mice relative to (PR)400 and WT mice (**Figure 3H**). These findings provide evidence for augmented neuronal and network excitability in motor cortex of (GR)400 mice.

### (GR)400 and (PR)400 knock-in mice develop age-dependent lower motor neuron loss and progressive rotarod impairment

Having identified early functional deficits in the brain, we next assessed motor function in our DPR knock-in mice. First, we measured body weight of (GR)400 and (PR)400 mice starting from 3-months of age. Our analysis did not detect differences in body weight over the course of the first year of life (**Fig. 4A and Supplementary Fig. 5A**). Similarly, we did not see alterations in body weight of eGFP knock-in mice from 3-to 12-months of age (**Supplementary Fig. 6A**). Next, we evaluated rotarod performance to establish whether polyGR or polyPR expression influenced motor coordination. Notably, we found a progressive decrease in accelerated rotarod performance (**Fig. 4B and Supplementary Fig. 5B**). In particular, (GR)400 mice showed significant rotarod impairment from 5-to 8-months of age, while in (PR)400 mice we found an impairment from 6-to 8-months of age when compared to WT littermates, which showed an age-related decline from 8 months of age onwards. As expected, we did not observe rotarod deficits in eGFP knock-in mice over the first year of life (**Supplementary Fig. 6B**), indicating the rotarod defect is a specific effect of DPR expression. We conducted grip strength analysis and neither (GR)400, (PR)400, nor eGFP knock-in mice showed strength deficits up to 12 months of age (**Supplementary Fig. 5C-D and Supplementary Fig. 6C**). These results show that polyGR and polyPR cause progressive, but relatively subtle motor dysfunction, consistent with physiological expression levels of the DPRs.

**Figure 4.**
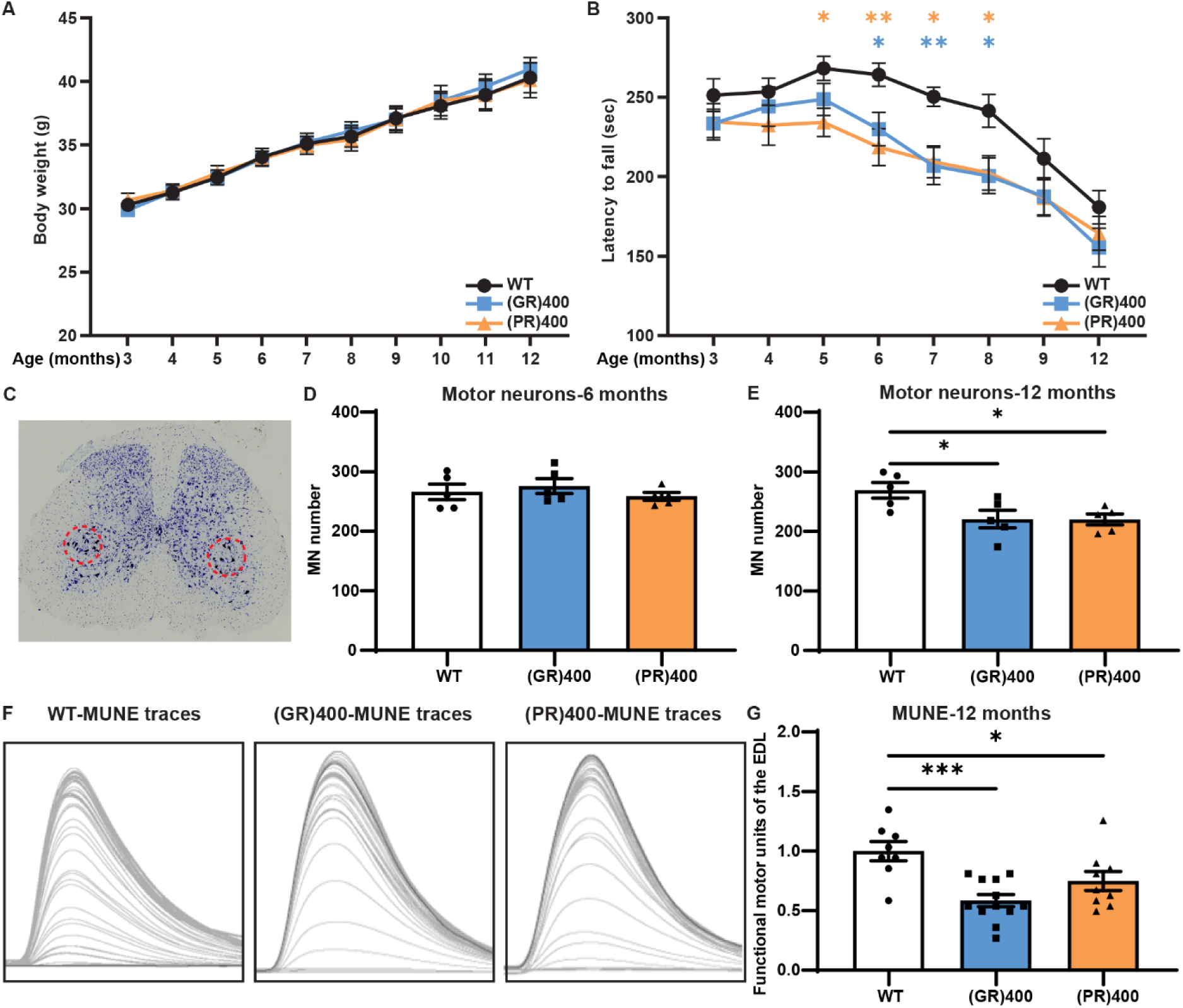
(GR)400 and (PR)400 knock-in mice develop age-dependent lower motor neuron loss and progressive rotarod impairment (A) Body weights of WT, (GR)400, and (PR)400 mice up to 12 months of age. Graph, mean ± SEM, n = 14 mice per genotype, two-way ANOVA, Bonferroni’s multiple comparison. (B) Accelerated rotarod analysis of motor coordination in WT, (GR)400, and (PR)400 mice up to 12 months of age. Graph, mean ± SEM, n = 14 mice per genotype, two-way ANOVA, Bonferroni’s multiple comparison, *p < 0.05, **p < 0.01. (C) Panel shows representative image of Nissl staining of lumbar spinal cord in WT mice; the dashed red line delineates the sciatic motor pool in which motor neurons were counted. (D) Quantification of Nissl-stained motor neurons in lumbar spinal cord region L3-L5 in WT, (GR)400, and (PR)400 mice at 6 months of age. Graph, mean ± SEM, n = 5 mice per genotype, one-way ANOVA, Bonferroni’s multiple comparison. (E) Quantification of Nissl-stained motor neurons in lumbar spinal cord region L3-L5 in WT, (GR)400, and (PR)400 mice at 12 months of age. Graph, mean ± SEM, n = 5 mice per genotype, one-way ANOVA, Bonferroni’s multiple comparison, *p < 0.05. (F) Representative EDL motor unit number estimation (MUNE) traces from 12-month old WT (left panel), (GR)400 (centre panel), and (PR)400 (right panel) mice. Peaks correspond to physiological recruitment of motor units following electrical stimulation. (G) Quantification of MUNE determined in EDL muscle in WT, (GR)400, and (PR)400 mice at 12 months of age. Graph, mean ± SEM, n mice = 5 WT, 6 (GR)400, and 6 (PR)400, n muscles = 8 WT, 12 (GR)400, and 9 (PR)400, one-way ANOVA, Bonferroni’s multiple comparison, *p < 0.05, ***p < 0.001.

We next turned our attention to neuropathological analysis of the ventral spinal cord, a key site of neurodegeneration in ALS. We focused on the ventral horn of the lumbar spinal cord as this is where the large alpha motor neurons that undergo degeneration in ALS reside. Similarly to the brain, we did not observe astrogliosis or microgliosis in the ventral horn of lumbar spinal cord from 12-month old (GR)400 and (PR)400 mice (**Supplementary Fig. 3B and 3D**). Moreover, no signs of TDP-43 cytoplasmic mis-localisation or aggregation were identified (**Supplementary Fig. 3F**).

Next, we determined whether polyGR or polyPR expression had an effect on lower motor neuron viability. We assessed neurodegeneration by counting motor neurons in the lumbar spinal cord (**Fig. 4C**). We did not observe any difference in the motor neuron number in 6-month-old (GR)400 and (PR)400 compared with WT mice (**Fig. 4D**). However, at 12 months of age, polyGR and polyPR mice showed a nearly 20% reduction in the number of motor neurons of the lumbar spinal cord (**Fig. 4E**). This is an important result as it shows that our knock-in mice replicate a cardinal feature of ALS: age-dependent spinal cord motor neuron loss. Given the importance of these findings, we next used *in vivo* electrophysiological recordings to validate loss of motor neurons. We performed motor unit number estimation analysis in the hind limbs of 12-month-old (GR)400 and (PR)400 mice. We detected a significant reduction, by 41% in (GR)400 and 25% in (PR)400, in functional motor unit number in the extensor digitorum longus (EDL) muscles when compared to wild-type littermates (**Fig. 4F and 4G**). Furthermore, we performed the same recordings in the hind limbs of 12-month old *C9orf72* eGFP knock-in mice without finding alterations in functional motor unit number (**Supplementary Fig. 6D**). Overall, these results reveal an age-dependent neurodegeneration in the spinal cord of our polyGR-and polyPR knock-in mouse models.

### (GR)400 and (PR)400 knock-in mouse spinal cord has increased extracellular matrix protein levels, a signature conserved in *C9orf72* patient motor neurons

Given that (GR)400 and (PR)400 mice showed features typical of ALS, we next investigated dysregulated pathways associated with DPR-induced toxicity using quantitative proteomics. Analysis of lumbar spinal cord of 12-month old (GR)400 and (PR)400 mice revealed a striking increase in extracellular matrix (ECM) terms (**Fig. 5A**), with no other clearly dysregulated pathways. Moreover, we obtained proteomic data from NeuroLINCS, which recently developed an integrated multi-omic analysis of C9ALS patient iPSC-derived motor neurons ^49^. We analysed their raw mass spectrometry data using our own analysis pipeline to ensure an appropriate comparison with our knock-in mouse data and observed a remarkable similarity between datasets, which showed a common upregulation of GO terms associated with the ECM, consistent with the original NeuroLINCS findings ^49, 50^ (**Fig. 5A**). Multiple ECM-associated proteins, including collagens, were pinpointed as the most significantly upregulated proteins in lumbar spinal cord of (GR)400 and (PR)400 mice, and iPSC-derived MNs (**Fig. 5B**). We also analysed a published dataset of laser-capture microdissected motor neurons from C9ALS patient spinal cord ^51^ and again extracellular matrix terms were among the most significantly upregulated GO terms (**Supplementary Fig. 7A**), consistent with a previous report ^50^. Fewer proteins were downregulated and with lower fold-changes than upregulated proteins, but interestingly, GO term enrichment analyses revealed synapse proteins were reduced in lumbar spinal cord of (GR)400 and (PR)400 mice (**Supplementary Fig. 7B**), consistent with the observed motor neuron loss. Importantly, a significant reduction in C9orf72 protein was observed, which is consistent with our quantitative PCR and immunoblotting data (**Fig. 5B**), helping confirm the quality of the dataset.

**Figure 5.**
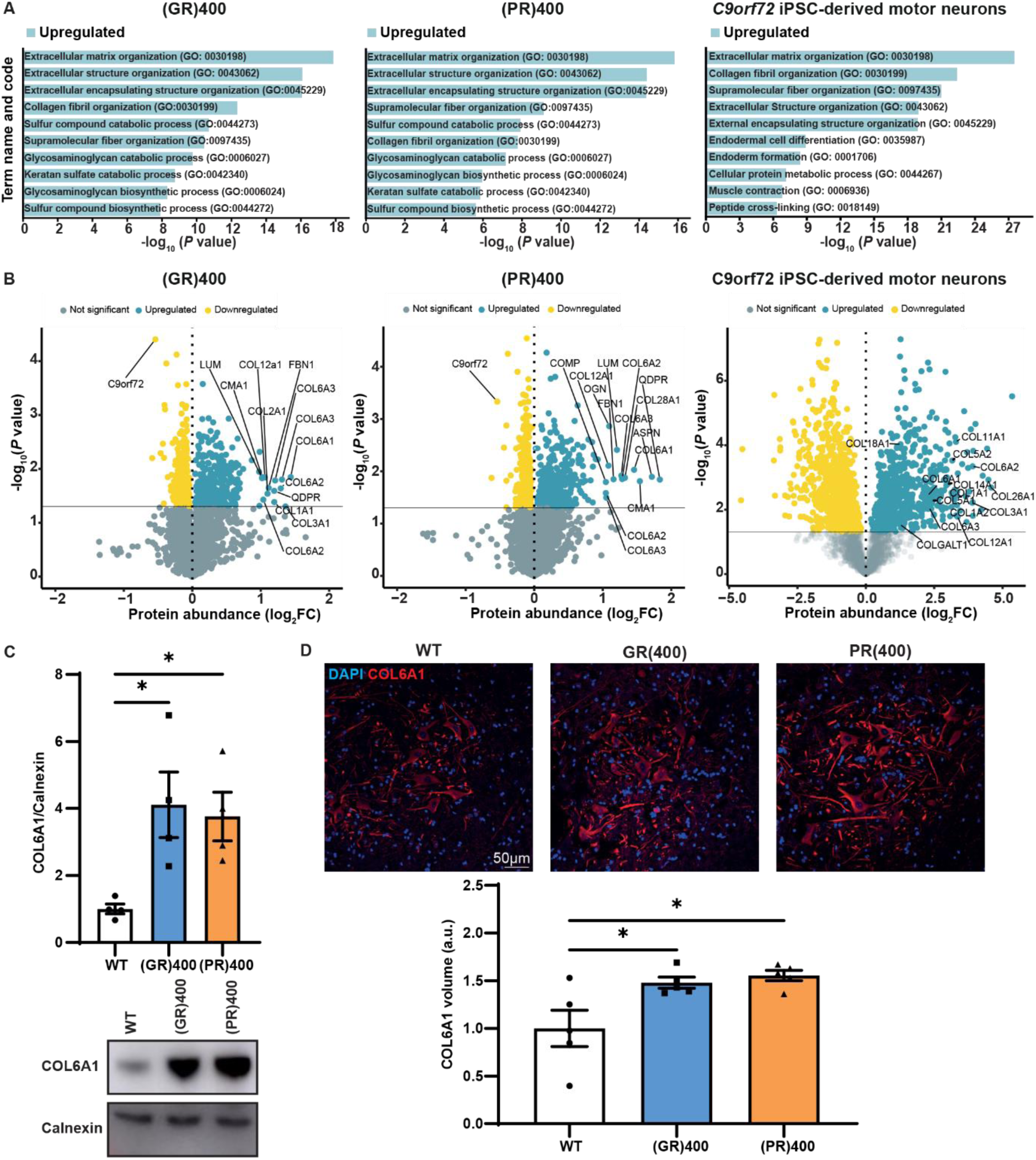
(GR)400 and (PR)400 knock-in mouse spinal cord has increased extracellular matrix protein levels (A) Gene Ontology (GO) term enrichment analysis from significantly upregulated (blue) proteins in the lumbar spinal cord of (GR)400 (left panel), or (PR)400 (centre panel), and *C9orf72* patient iPSC-derived motor neurons (right panel). Proteomics performed on n = 5 mice per genotype. (B) Protein expression volcano plots from the lumbar spinal cord of (GR)400 (left panel), (PR)400 (centre panel), and *C9orf72* patient iPSC-derived motor neurons (right panel). (C) Western blot of COL6A1 in lumbar spinal cord of WT, (GR)400, and (PR)400 mice at 12 months of age. Calnexin (CNX) is shown as loading control. Graph, mean ± SEM, n = 4 mice per genotype, one-way ANOVA, Bonferroni’s multiple comparison, *p < 0.05. (D) Representative confocal images and quantification of immunofluorescence staining of COL6A1 (red) in lumbar spinal cord in WT, (GR)400, and (PR)400 mice at 12 months of age. DAPI (blue) stains nuclei. Graph, mean ± SEM, n = 5 mice per genotype, one-way ANOVA, Bonferroni’s multiple comparison, *p < 0.05.

To further verify our results, we evaluated COL6A1 expression in (GR)400 and (PR)400 mice as it was one of the most altered ECM proteins in the proteomics analysis. As expected, immunoblot analysis showed COL6A1 was significantly upregulated in lumbar spinal cord of 12-month old (GR)400 and (PR)400 mice (**Fig. 5C**). Similarly, immunostaining followed by volumetric image analysis showed a ∼50% increase of COL6A1 volume in the ventral horn of lumbar spinal cord from (GR)400 and (PR)400 mice at 12-months of age (**Fig. 5D**), but not eGFP knock-in mice (**Supplementary Fig. 7C**). Intriguingly, COL6A1 was localised to neurons rather than the extracellular space or other cell types, indicating an increase of neuronal ECM protein expression. This is consistent with the increased ECM signature in patient iPSC-motor neurons and laser capture microdissected motor neurons. Overall, these data show that an increase in extracellular matrix proteins, exemplified by COL6A1, is a conserved feature of *C9orf72* FTD/ALS neurons.

### PolyGR induces TGF-β1 followed by its target gene *COL6A1* in i^3^Neurons

To identify potential upstream regulators controlling the differential expression of ECM-related proteins, we conducted Ingenuity Pathway Analysis (IPA) on the polyGR, polyPR and patient iPSC-motor neuron data sets. IPA uncovered several predicted regulators. Several of the top predicted regulators were common between our DPR knock-in mice and human *C9orf72* iPSC-derived motor neurons, including TGF-β1 and its intracellular mediator SMAD2/3, as well as AGT, CCR2 and SORL1 (**Fig. 6A**). Interestingly, among these regulators, we found that TGF-β1 was significantly upregulated within the 3 proteomic datasets (**Supplementary Table 1 and Supplementary Table 2**). To determine whether TGF-β1 is also altered in patient brain we utilised a large frontal cortex RNA-seq dataset comprising 34 *C9orf72* FTD/ALS cases in which ECM dysregulation has also been reported ^52^. We found that *TGFB1* was significantly increased, after genome-wide FDR correction, in *C9orf72* FTD/ALS cases when compared to either non-*C9orf72* FTD/ALS or neurologically normal controls (**Fig 6B**), further confirming the relevance of this pathway in *C9orf72* FTD/ALS patient tissue.

**Figure 6.**
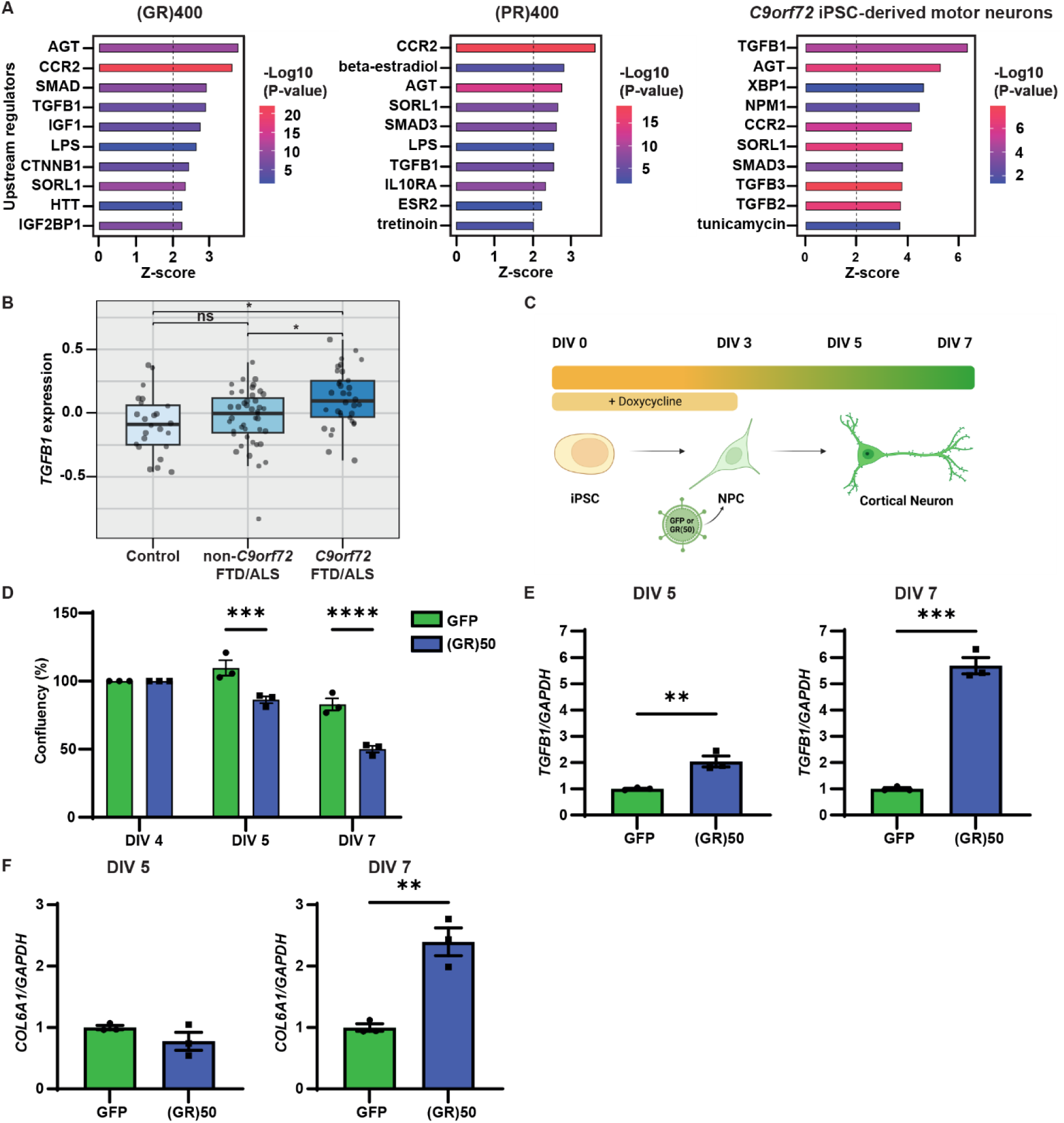
PolyGR induces TGF-β1 followed by its target gene COL6A1 in i^3^Neurons (A) The top 10 IPA-predicted upstream regulators in (GR)400 (left panel), (PR)400 (centre panel) and *C9orf72* iPSC-derived motor neurons (right panel). B) Boxplot of residual gene expression for *TGFB1*. The solid black line represents the median and the box represents the interquartile range (IQR; 25th percentile to 75th percentile). Significance was determined by differential gene expression analysis using a linear regression model. P-values are FDR corrected, accounting for all genes included in the differential expression analysis. * FDR < 0.05; ns FDR > 0.05. *C9orf72* FTLD/MND n = 34, non-*C9orf72* FTLD/MND n = 44, controls n = 24. (C) Schematic workflow of i^3^Neuron doxycycline-induced differentiation at DIV 0 and transduction with lentiviruses expressing (GR)_50_, or GFP at DIV 3. (D) Live-cell Incucyte confluency quantification in (GR)_50_, or GFP-treated i^3^Neurons. Graph, mean ± SEM, n = 3 independent biological replicates, two-way ANOVA, Bonferroni’s multiple comparison, ***p < 0.001, ****p < 0.0001. (E) Quantitative PCR analysis of *TGFB1* transcript levels normalised to *GAPDH* at DIV 5 (left panel) and DIV 7 (right panel) in (GR)_50_, or GFP-treated i^3^Neurons. Graph, mean ± SEM, n = 3 independent biological replicates, one-way ANOVA, Bonferroni’s multiple comparison, **p < 0.01, ***p < 0.001. (F) Quantitative PCR analysis of *COL6A1* transcript levels normalised to *GAPDH* at DIV 5 (left panel) and DIV 7(right panel) in (GR)_50_, or GFP-treated i^3^Neurons. Graph, mean ± SEM, n = 3 independent biological replicates, one-way ANOVA, Bonferroni’s multiple comparison, **p < 0.01.

Based on these results, we hypothesized that activation of the TGF-β1 signalling pathway, which is known to be a master regulator of ECM genes ^53^, may contribute to the ECM alterations in (GR)400 and (PR)400 mice. To test this hypothesis, we studied TGF-β1 signalling *in vitro*. We adopted a transcription factor-mediated differentiation protocol to differentiate human induced pluripotent stem cells into cortical neurons (i^3^Neurons) ^54^. We transduced i^3^Neurons with 50 polyGR repeats [(GR)_50_] ^55^, or GFP as a negative control (**Fig. 6C**). (GR)_50_ caused a progressive increase in neuronal death, with a –20% and ∼40% reduction in confluency at 5 and 7 days *in vitro* (DIV) respectively compared to GFP-treated cells (**Fig. 6D**). Quantitative PCR analysis showed that *TGFB1* was significantly upregulated by polyGR at DIV 5, prior to its target gene *COL6A1*, which became significantly increased two days later (**Fig 6E and 6F**). This shows that polyGR expression in neurons is sufficient to induce TGF-β1 expression, leading to increased expression of *COL6A1*, the most prominently increased ECM protein in polyGR knock-in mouse spinal cord. This suggests that increased TGF-β1 signalling is one factor contributing to the conserved ECM protein signature in C9 FTD/ALS neurons.

### TGF-β1 and COL6A1 reduction specifically exacerbates polyGR toxicity *in vivo*

The neuronal increase in ECM proteins could be protective, deleterious or neutral to disease progression. To investigate this further we focused on TGF-β1, as it emerged as a top predicted regulator, and COL6A1, as it is the most highly increased ECM protein in polyGR mice. To determine whether *TGF-β1* and *COL6A1* have a role in polyGR-induced neurodegeneration *in vivo*, we utilised our *Drosophila* line expressing 36 polyGR repeats ^15, 56^, which causes a moderate eye degeneration. We investigated the *dawdle* (*daw*) and *Multiplexin* (*Mp*) genes, which are the closest fly orthologues of *TGFB1* and *COL6A1*, respectively. Both *daw* and *Mp* levels were increased in GR36 flies (**Fig 7A**), consistent with both the knock-in mouse and patient iPSC-motor neuron data. We crossed GR36 flies with RNAi lines targeting *daw* or *Mp*. Both *daw* and *Mp* reduction caused a dramatic worsening of eye degeneration in GR36 flies but had no effect in wildtype flies (**Fig. 7B-7D**). This shows that *daw* and *Mp* are able to specifically protect against polyGR-induced toxicity *in vivo*, indicating that increased neuronal collagen expression is neuroprotective in the context of polyGR insult. In summary, ECM proteins, exemplified by COL6A1 are specifically increased in the spinal cord of our new DPR knock-in mice, as well as *Drosophila* and human iPSC-neurons expressing polyGR, patient iPSC-motor neurons and patient end-stage spinal-motor neurons. The presence of this conserved signature in surviving neurons across different models and patient material, combined with our data showing a protective role in GR36 *Drosophila* indicates a novel neuroprotective role of neuronally expressed ECM proteins.

**Figure 7.**
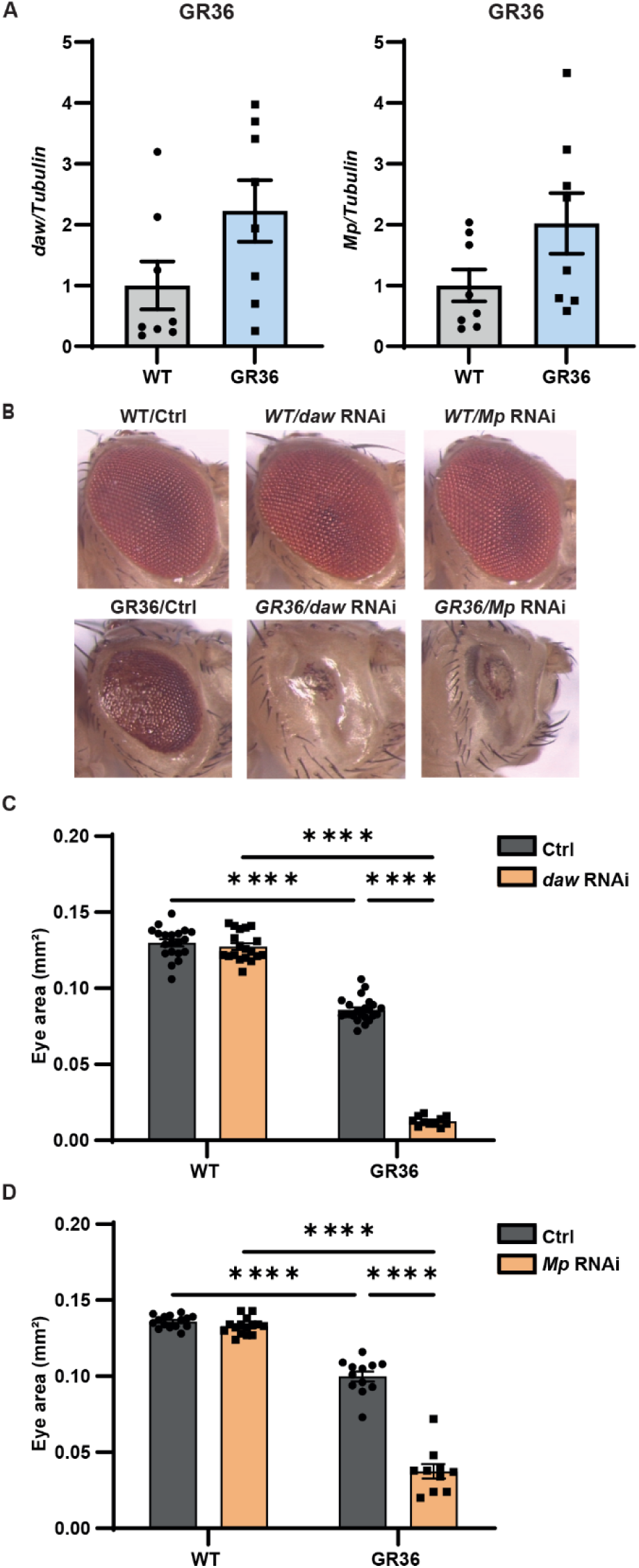
TGF-β1 and COL6A1 reduction specifically exacerbates polyGR toxicity *in vivo* (A) qPCR analysis of *daw* (left panel) and *Mp* (right panel) transcript levels normalised to *Tubulin* in WT and GR36 flies. Graph, mean ± SEM, n = 8 independent biological replicates, unpaired two-sample Student’s t-test. Genotypes: w; GMR-Gal4/+, w; GMR-Gal4, UAS-GR36/+. (B) Stereomicroscopy images of representative 2-day old adult WT (top panel) or GR36 (bottom panel) *Drosophila* eyes in absence or co-expressing *daw* or *Mp* RNAi constructs. Genotypes: w; GMR-Gal4/+, w; GMR-GAL4/UAS-*daw* RNAi, w; GMR-GAL4/UAS-*Mp* RNAi, w; GMR-Gal4, UAS-GR36/+, w; GMR-Gal4, UAS-GR36/UAS-*daw* RNAi, w; GMR-Gal4, UAS-GR36/UAS-*Mp* RNAi. (C) Eye size of flies normalized to the mean of the control eye size. Graph, mean ± SEM, n (independent biological replicates, one eye counted per fly) = 20 WT/+, 19 WT/*daw* RNAi, 23 GR36/+, 11 GR36/*daw* RNAi, two-way ANOVA, Bonferroni’s multiple comparison, ****p < 0.0001. Genotypes: w; GMR-Gal4/+, w; GMR-GAL4/UAS-*daw* RNAi, w; GMR-Gal4, UAS-GR36/+, w; GMR-Gal4, UAS-GR36/UAS-*daw* RNAi. (D) Eye size of flies normalized to the mean of the control eye size. Graph, mean ± SEM, n (independent biological replicates, one eye counted per fly) = 15 WT/+, 15 WT/*Mp* RNAi, 12 GR36/+, 10 GR36/*Mp* RNAi, two-way ANOVA, Bonferroni’s multiple comparison, ****p < 0.0001. Genotypes: w; GMR-Gal4/+, w; GMR-GAL4/UAS-*Mp* RNAi, w; GMR-Gal4, UAS-GR36/+, w; GMR-Gal4, UAS-GR36/UAS-*Mp* RNAi.

## Discussion

We have generated the first DPR knock-in mice, which utilise the endogenous mouse *C9orf72* promoter to drive expression of a single DPR, either (GR)400 or (PR)400. The insertion of the DPR sequence also removed one normal allele of *C9orf72.* Thereby we have recapitulated two features of *C9orf72* FTD/ALS – the reduced level of C9orf72 and the presence of DPRs. The mice do not recapitulate the unconventional RAN translation mechanism of DPR generation, as they are driven by the mouse *C9orf72* ATG start codon. This was intentional, allowing us to study the effects of specific DPRs, and means that other mouse models, expressing the expanded G_4_C_2_ repeat, are needed to study mechanisms of RAN translation *in vivo*. We focussed on polyGR and polyPR as they are consistently the most toxic DPRs in model systems ^29^. This does not lessen the relevance of other DPRs, particularly polyGA, which is also toxic in different model systems, but likely through different mechanisms ^14, 57^.

We show that driving expression with the endogenous mouse promoter leads to physiological expression levels of polyGR, as determined by comparison with human brain, allowing the examination of patient-relevant polyGR effects. We also built DPR constructs in the patient range, comprising 400 uninterrupted polyGR or polyPR repeats, to replicate patient DPRs as closely as possible. Physiological expression of patient-length repeats did not lead to overt gliosis or TDP-43 mis-localisation over 12 months, nor did it lead to cortical neuronal loss. This is consistent with the milder effects observed in knock-in mice, in which genes are expressed at physiological levels, and provides reassurance that over-expression artefacts are unlikely ^44^. Over-expression of 200 GR repeats, but not 100 GR repeats was sufficient to drive aggregation of polyGR and recruitment of TDP-43 into those aggregates, implicating GR aggregation as a possible cause of TDP-43 aggregation ^24^. Our expression of even longer repeats, but at lower levels did not lead to GR or TDP-43 aggregation, suggesting a complex relationship between DPRs and TDP-43 aggregation.

Intriguingly, using two complementary methods, namely two-photon calcium imaging and high-density Neuropixels recordings, we observed hyperexcitability in polyGR mice in superficial cortical layers, which are affected in FTD, but not deeper layer 5 neurons that are vulnerable in ALS. Further work is needed to parse out this difference and its relevance, but it is noteworthy that cortical hyperexcitability is well described in both *C9orf72* and non-*C9orf72* ALS/FTD patients ^58^, indicating that new mechanistic insights into this early patient-relevant phenotype can be gained by future investigations on our polyGR mice. It is also interesting that only polyGR but not polyPR mice showed this phenotype. Firstly, this rules out the effect being due to reduced C9orf72 levels, as both mice have similarly reduced C9orf72. Secondly, it points to a more pertinent effect of polyGR in the brain. This would be consistent with several studies that show polyGR is the only DPR to correlate with clinical symptoms and neurodegeneration in C9FTD/ALS patients ^59–61^. PR20 peptides were shown to cause cortical neuron excitability in acute brain slices ^62^, so an effect of polyPR cannot be ruled out, but our use of *in vivo* recording and patient-length repeats appears closer to the patient context.

It was also notable that polyGR expression appeared restricted to neurons when driven by the mouse *C9orf72* promoter. This is consistent with two earlier reports that drove LacZ with the mouse *C9orf72* promoter ^63, 64^, as well as *in situ* hybridisation and immunostaining of mouse *C9orf72*, which showed only neuronal and no glial staining in mouse brain ^65^. This is also consistent with *C9orf72* repeat RNA foci and DPRs being expressed at lower levels in glia than neurons ^8, 10, 60, 66, 67^. However, as *C9orf72* knockout microglia exhibit an altered transcriptome and functional properties ^68^, further investigation of the circumstances under which a microglial function for C9orf72 becomes apparent are warranted.

While we did not observe neuronal loss in the cortex, we did observe a significant, approximately 20% loss of spinal cord motor neurons in both polyGR and polyPR mice. We therefore focused our mechanistic analyses on the spinal cord. A striking finding was a remarkably conserved signature of increased ECM gene expression in *C9orf72* FTD/ALS tissues. The increased ECM signature was the dominant change in quantitative proteomics analyses of both polyGR and polyPR knock-in spinal cord and it was remarkably similar to an independent quantitative proteomics dataset of *C9orf72* patient iPSC-motor neurons. This increase was also present in laser capture micro-dissected *C9orf72* patient spinal cord motor neurons, human iPSC-neurons treated with polyGR and *Drosophila* over-expressing polyGR. This conservation across several models and human tissue indicates a genuine phenomenon linked to the presence of arginine-rich DPRs. Altered ECM gene expression has been noted in range of ALS-related transcriptomic datasets both with and without *C9orf72* mutation, but there is no clear consensus on its role and importance in ALS pathophysiology. It is clear that one driver of increased ECM gene expression in those datasets is astrogliosis. A meta-analysis of human iPSC-astrocyte transcriptomic datasets from several genetic subtypes of ALS (*C9orf72*, *FUS*, *SOD1* and *VCP*) revealed a common increase in ECM gene expression that was shared with pro-inflammatory ‘A1’ astrocytes ^69^. This is consistent with recently published bulk spinal cord RNA-seq data from a large series of 154 ALS cases (including 29 *C9orf72* cases ^70^). Weighted gene co-expression network analysis of this dataset identified 23 co-expressed gene modules, including an astrocyte module with significantly increased ECM gene expression in ALS cases that was negatively correlated with age at onset and age at death ^70^. This suggests that pro-inflammatory astrocytes are characterised by increased ECM gene expression and that this correlates with more severe disease. Interestingly, ALS microglia also appear to show increased ECM gene expression ^70^ indicating ECM signals in bulk transcriptomic data could derive from both activated astrocytes and microglia, as well as neurons. Here we show a neuron-centric alteration in the absence of overt gliosis; however, these studies warrant future careful examination of the role of glia in ECM changes in *C9orf72* FTD/ALS.

Our data now brings clarity to the complexities of ECM alterations in ALS/FTD by showing that there is a neuron-derived ECM signature that is neuroprotective. This provides a new paradigm for understanding ECM changes in which a potentially deleterious ECM increase in glia occurs alongside a protective ECM upregulation in neurons. These opposing effects may help explain why it has previously been hard to pinpoint the role and relevance of ECM alterations in ALS/FTD. The case for neuron-derived aberrant ECM expression is supported by its presence in neuronal datasets – patient iPSC-motor neurons and laser capture microdissected motor neurons, as well increased COL6A1 immunostaining in motor neurons in polyGR and polyPR knock-in mice. This ECM signature appears to be driven, at least in part, by TGF-β1, as TGF-β1 itself is increased and it was one of the top predicted upstream regulators of the changes in polyGR, polyPR and patient iPSC-neuron proteomics datasets. It is possible that the other predicted regulators also contribute to the increased ECM signal and this will require further investigation. TGF-β1 is an attractive candidate as it is well known to cause increases in ECM expression via its intracellular mediator phospho-SMAD2/3 ^53^, and expression of polyGR was sufficient to induce *TGFB* expression, providing a plausible explanation for its upregulation. In addition, TGF-β1 is neuroprotective both *in vitro* and *in vivo* against a wide array of neuronal insults including excitotoxicity ^71–73^, hypoxia ^74^ and ischemia ^75, 76^ and *Tgfb* knockout mice exhibit neurodegeneration ^77, 78^. We now show TGF-β1 expression can lead to a neuroprotective increase in the neuronal expression of ECM proteins and specifically collagen VI.

Expression of polyGR in human iPSC-neurons was sufficient to induce *TGFB1* expression followed by *COL6A1*, and knockdown of the *COL6A1* or *TGFB1* homologues in flies expressing 36 GR repeats showed a specific and striking enhancement of neurodegeneration, pointing to a neuroprotective effect of collagen induction. In remarkable concordance with our data, it was previously shown that Aβ_42_ treatment of primary mouse neurons causes an increase in neuronal Col6a1 that is neuroprotective and mediated by TGF-β1 ^79^. Neuronal collagen VI expression is also increased upon UV-irradiation of primary neurons and protects against irradiation-induced apoptosis ^80^. In combination, these results show that *COL6A1*, which is not normally expressed in neurons, is induced in neurons by neurodegenerative insults. Collagen VI is comprised of a 1:1:1 polymer of three distinct collagen VI chains encoded by *COL6A1*, *COL6A2* and *COL6A3*. This heterotrimer then tetramerises before being secreted into the extracellular space where it classically forms beaded microfilaments ^81^. However, in neuronal cultures collagen VI was identified in proximity to the neuronal plasma membrane indicating that it may act as a neuroprotective autocrine signalling molecule when secreted by neurons rather than forming microfilaments ^82^. This is supported by experiments in which exogenously added collagen VI was sufficient to ameliorate both UV-irradiation and Aβ_42_-induced neuronal death ^79, 80^. However, further investigations are now needed to identify the mechanism by which collagen VI provides neuroprotection. All three collagen VI chains were upregulated in our proteomics datasets so it will be important to determine whether individual collagen VI chains such as COL6A1 can perform independent neuroprotective functions or whether the intact final tetramer is the active species.

It is clear that a range of different insults can induce TGF-β1/ECM changes. One such insult is polyGR/polyPR, but it is likely that other ALS-related insults can also lead to TGF-β1 and collagen increase. Indeed, dysregulated TGF-β1 signalling has been described in a *SOD1* mouse model of ALS ^83–85^ and increased ECM and TGF-β1 signalling was identified in a transcriptomic dataset of sporadic ALS post-mortem spinal motor neurons ^50^. This indicates that different insults relevant to genetic or sporadic forms of ALS can cause a TGF-β1 response. Some studies report that TGF-β1 concentration is significantly higher in plasma ^86^ and cerebrospinal fluid ^87^ from ALS patients compared to healthy controls. Although other studies have not observed these alterations in ALS patients ^88^. It is therefore still unknown whether TGF-β1 biofluid levels may be a suitable biomarker for ALS ^89^. In summary, these findings identify a neuroprotective neuronal ECM signature in *C9orf72* ALS/FTD, exemplified by collagen VI, which may have broad relevance for ALS and other neurodegenerative diseases.

## Methods

### Assembly of targeting constructs

eGFP sequence and 100 codon-optimised polyGR or polyPR were synthesized at GeneArt Synthesis Services (Thermo Fischer Scientific). An Empty Vector (EV) was generated at GeneArt Synthesis Services (Thermo Fischer Scientific) by engineering a pMC cloning vector to contain 600 bp of the 5’ homology arm, a double HA-tag, a V5 epitope tag, a SV40 polyA tail, and 250 bp of 3’ homology arm. eGFP and 100 codon-optimised DPRs were cloned into the EV within the *BbsI* and *BsmBI* sites to generate pMC-eGFP, pMC-(GR)100, and pMC-(PR)100. 400 codon-optimised DPRs were assembled with two consecutive rounds of recursive directional ligation taking advantage of the restriction enzymes *BbsI* and *BsmBI* to generate pMC-(GR)400 and pMC-(PR)400 A selection cassette (FRT-PGK-gb2-neo-FRT, Gene Bridges) was inserted in the pMC-eGFP, pMC-(GR)400, and pMC-(PR)400 in the *NheI* site. A BAC subcloning kit by Red/ET recombination (Gene Bridges) was used to clone full length homology arms (2.7 kb in 5’ and 3.2 kb in 3’) from the BAC clone (RP23-434N2), containing the C57BL/6J sequence of the mouse *C9orf72* gene, into the targeting vector [pBlueScript II SK (+)]. Knock-in constructs were obtained by inserting sequences from pMC-eGFP, pMC-(GR)400, and pMC-(PR)400 into targeting vectors within the *BstXI* and *XcmI* sites.

### Animals

All procedures involving mice were conducted in accordance with the Animal (Scientific procedures) Act 1986 and performed at University College London under an approved UK Home Office project licence. Mice were maintained in a 12 h light/dark cycle with food and water supplied *ad libitum*.

To generate the KI-DPR mouse strains, we performed CRISPR assisted gene targeting in JM8F6 embryonic stem (ES) cells (C57BL/6N) using our targeting vector(s) and a CRISPR/Cas9 designed against the insertion site.

The CRISPR construct (pX330-Puro-C9orf72), expressing Cas9 and a U6 promoter driven single-guide-RNA (sgRNA) designed against the following sequence AGTCGACATCCCTGCATCCC. This was generated by annealing the following two oligos (5’-CACCgAGTCGACATCCCTGCATCCC-3’; 5’-AAACGGGATGCAGGGATGTCGACTc-3’) and cloning this into the unique *BbsI* sites of pX330-U6-Chimeric_BB-CBh-hSpCas9 (Addgene # 42230), modified by the addition of a PGK-Puro cassette.

1×10^6^ ES cells were electroporated with 2.5 µg of the cloned pX330-Puro-C9orf72 plasmid and 2.5 µg of targeting vector using the Neon Transfection System (Thermo Fisher Scientific) (3 × 1400 V, 10 ms) and plated on Puromycin resistant fibroblast feeder layers. After approximately 24 h, selection in 600 ng/ml puromycin was applied for a further 48 h to allow transient selection. After further 5 days in culture without selection, individual colonies were isolated, expanded and screened for the desired targeting at both the 5’ end (5’-TCGGGGATTATGCCTGCTGC’3’ and 5’-GCATCCCAGGTCTCACTGCA-3’) and at the 3’ end (5’-TCGAAAGGCCCGGAGATGAGGAAG-3’ and 5’-GGGTTCAGACAGGTACAGCAT-3’). ES cells from correctly targeted clones were injected into albino C57BL/6J blastocysts and the resulting chimeras were mated with albino C57BL/6J females. The presence of the targeted allele in the F1 generation was confirmed at the DNA level by the above PCR and Sanger sequencing. Germline-transmitting founders were obtained and backcrossed to wild-type C57BL/6J mice to maintain hemizygous lines.

Mouse genotype was determined by PCR for knock-in sequence with the following set of primers (forward 5’-TAAGCACAGCAGTCATTGGA-3’ and reverse 5’-AAGCGTAATCTGGAACATCG-3’). Repeat length was determined by PCR with the following set of primers (forward 5’-CCCATACGATGTTCCAGATTACGCTTACCC-3’ and reverse 5’-GCAATAAACAATTAGGTGCTATCCAGGCCCAG-3’).

Males were used for all experiments in the main text, except *in vivo* two-photon calcium imaging and Neuropixels recording where females were used. Phenotyping was also performed on female mice, with similar results to males, and these data are included in the supplement data section.

### Human tissues

Human C9orf72FTD/ALS samples for polyGR MSD immunoassay were described previously ^61^ and protein extracted as described in the biochemical analysis section.

### Biochemical analysis

Brains and spinal cords were mechanically pulverized and homogenized in lysis buffer [RIPA buffer (Pierce), 2% sodium dodecyl buffer (SDS)] containing protease (Roche) and phosphatase (Thermo Scientific) inhibitor cocktails. The lysates were sonicated and microcentrifuged for 20 min at 13000 rpm at room temperature. Soluble fractions were collected and protein concentration was determined using the bicinchoninic acid (BCA) assay method (Thermo Scientific). Protein were separated on NuPAGE™ 4 to 12% bis-tris gels (Invitrogen) andtransferred to nitrocellulose membranes (Bio-Rad Laboratories). Membranes were blocked in 5% milk in PBS-T (PBS, 0.1% Tween-20) for 1 hour at room temperature. The membranes were incubated overnight at 4 °C with the following primary antibodies: C9orf72 (12E7, kindly donated by Prof. Dr. Manuela Neumann; 1:4 dilution), COL6 (ab182744, Abcam; 1:1000), β-Actin (A2228, Sigma-Aldrich;1:5000 dilution), Calnexin (sc-6465, Santa Cruz Biotechnology; 1:1000 dilution). After 3 washes in PBS-T, membranes were incubated with secondary HRP-conjugated antibodies for 1 h at room temperature. After 3 washes in PBS-T, signals were visualized by chemiluminescence (Amersham imager 680) and quantifications were performed using ImageJ software.

For Meso Scale Discovery (MSD) immunoassay, 96-well MSD microplates were coated with a capture antibody in PBS and placed overnight at 4°C. Plates were washed 3 times with TBS-T (TBS, 0.2% Tween-20) and blocked with 3% milk in TBS-T with shaking for 2 hours at room temperature. Plates were washed 3 times with TBS-T, then samples and protein standards were loaded and left shaking overnight at 4°C. After 3 washes in TBS-T, plates were incubated with a biotinylated detector antibody and left shaking for 2 h at room temperature. After 3 washes in TBS-T, plates were incubated with streptavidin-sulfotag (R32AD1, Meso Scale Discovery) and left shaking for 1 h at room temperature. Then plates were washed 3 times in TBS-T and MSD reading buffer (R92TC-1, Meso Scale Discovery) was added to samples and protein standards. Plates were read with a MSD Discovery microplate reader at 620 nm. Capture antibodies were: our previously described custom rabbit anti-(GR)_7_ antibody (Eurogentec 2 µg/mL) ^90, 91^, PR32B3 (Helmholtz Zentrum 2 µg/mL). Detector antibodies were: the same GR antibdut used for capture (1 µg/mL, biotinylated), PR32B3 (2 µg/mL, biotinylated in house).

GFP levels were measured by the GFP ELISA Kit (ab171581, Abcam) according to manufacturer’s instructions.

### Quantitative reverse transcription PCR

Tissues were dissected and flash-frozen in isopentane precooled in dry-ice, and stored at – 80°C until further processing. Total RNA was extracted with miRNeasy Micro Kit (Qiagen) and reverse-transcribed into cDNA using SuperScript IV Reverse Transcriptase (Invitrogen) with random hexamers and Oligo(dT)_20_ Primer. Gene expression was determined by quantitative real-time PCR using LightCycler^®^ 96 (Roche). Relative gene expression was determined using ΔΔCT method. For quantitative PCR, primers specific for mouse *C9orf72* are: forward primer 5’-TGAGCTTCTACCTCCCACTT-3’ and reverse primer 5’-CTCTGTGCCTTCCAAGACAAT-3’. Primers for mouse *Actin* are: forward primer 5’-CTGGCTCCTAGCACCATGAAGAT-3’ and reverse primer 5’-GGTGGACAGTGAGGCCAGGAT-3’. These primers were used in reactions with LightCycler® 480 SYBR Green I Master (Roche).

Cells were collected, pelleted, and frozen at –80°C until further processing. Total RNA was extracted with ReliaPrep^TM^ RNA Cell Miniprep System (Promega) and reverse-transcribed into cDNA using SuperScript IV VILO Master Mix (Invitrogen). Gene expression was determined by quantitative real-time PCR using LightCycler^®^ 96 (Roche). Relative gene expression was determined using ΔΔCT method. For quantitative PCR, primers specific for human *TGFB1* are: forward primer 5’-GGCTACCATGCCAACTTCT-3’ and reverse primer 5’-CCGGGTTATGCTGGTTGT-3’. Primers for human *COL6A1* are: forward primer 5’-ACTTCGTCGTCAAGGTCATC-3’ and reverse primer 5’-CATCTGGCTGTGGCTGTA-3’. Primers for human *GAPDH* are: forward primer 5’-ACTAGGCGCTCACTGTTCT-3’ and reverse primer 5’-CCAATACGACCAAATCCGTTG-3’. These primers were used in reactions with LightCycler® 480 SYBR Green I Master (Roche).

*Drosophila* were flash-frozen in liquid nitrogen. Total RNA was isolated using TRIzol reagent (Thermo Fisher Scientific) following the manufacturer’s protocol. RNA samples were treated with TURBO DNase (Thermo Fisher Scientific), and converted to cDNA using oligod(T) primers and Superscript II reverse transcriptase (Invitrogen). Gene expression was determined by quantitative real-time PCR using QuantStudio 6 Flex Real-Time PCR System (Applied Biosystems). Relative gene expression was determined using ΔΔCT method. For quantitative PCR, primers for *Drosophila daw* are: forward primer 5’-GGATCAGCAGAAGGACTCCAA-3’ and reverse primer 5’-CAGTGTTTGATGGGCCACTC-3’. Primers for *Drosophila Mp* are: forward primer 5’-CTGGGCACCTTCAAGGCATT-3’ and reverse primer 5’-ATCGCCACGAGTGTTCACC-3’. Primers for *Drosophila Tubulin* are: forward primer 5’-TGGGCCCGTCTGGACCACAA-3’ and reverse primer 5’-TCGCCGTCACCGGAGTCCAT-3’. These primers were used in reactions with SYBR Green Master Mix (Applied Biosystems).

### Immunohistochemistry

Mice were anesthetized by isofluorane inhalation and perfused through a transthoracic cardiac puncture with prechilled phosphate buffered saline (PBS) and then 4% paraformaldehyde (PFA). Brains and spinal cords were dissected and postfixed in 4% PFA at 4°C for 2 h. After fixation, brains and spinal cords were washed with PBS, allowed to sink in 30% sucrose solution at 4°C, then stored in 0.02% sodium azide at 4°C until further processing. Brains and spinal cords were embedded in optimal cutting temperature (OCT) compound (Tissue Tek, Sakura, Torrance, CA), and cross sections (10 μm-thick) were cut with a cryostat (CM1860 UV, Leica Microsystem), then stored at –80°C. For immunofluorescence analysis, brains and spinal cord cryosections were washed 3 times in PBS and blocked in blocking solution (5% BSA, 1% normal goat serum, 0.2% Triton-X in PBS) for 1 h at room temperature. Sections were then incubated with primary antibodies in blocking solution overnight at 4°C. After 3 washes with PBS, sections were incubated for 1 hour at room temperature in blocking solution with secondary antibodies conjugated with Alexa 488, 546, 594, and 633 (Invitrogen). After 3 washes in PBS, sections were mounted with ProLong™ Gold Antifade Mountant with DAPI (Invitrogen).

Alternatively, after fixation, brains were washed with PBS, processed overnight using an automated tissue processor (Leica ASP300), and embedded in paraffin (Leica EG1150H). Tissue blocks were then stored at room temperature. For immunofluorescence analysis, 5 µm sections mounted on glass slides were incubated for 2 h at 60 °C. Sections were deparaffinised in xylene and rehydrated in decreasing grades of alcohol. Slides were incubated in methanol/hydrogen peroxide (0.3%) solution for 10 min at room temperature to block endogenous peroxidase activity. For heat-induced antigen retrieval, slides were then transferred to a boiling solution of 0.1 M citrate buffer (pH 6.0) and pressure cooked at maximum pressure for 10 min. For immunofluorescence, slides were then blocked in 10% milk for 1 hour at room temperature and incubated with primary and secondary antibodies as described above. For DAB staining, slides were incubated in methanol/hydrogen peroxide (0.3%) solution for 10 min at room temperature to block endogenous peroxidase activity, prior to antigen retrieval as described above. Slides were then blocked in 10% milk for 1 h at room temperature and incubated with primary antibody in PBS overnight at 4°C. After 3 washes with PBS, sections were incubated for 30 min at room temperature in biotinylated secondary antibody (Vector Laboratories) in PBS. Slides were then washed in PBS and incubated in VECTASTAIN® Elite® ABC-HRP Kit, Peroxidase (Vector Laboratories) for 30 min at room temperature. Sections were washed 3 times with PBS and incubated in 3,3ʹ-Diaminobenzidine (DAB) chromogen (Abcam).Slides were then dehydrated in increasing grades of alcohol (70%, 95% and 100% ethanol), cleared in xylene and mounted with DPX mounting medium (Sigma-Aldrich).

The primary antibodies used were: HA clone 3F10 (11867423001, Roche; 1:100 dilution), NEUN (ABN91, Millipore; 1:500 dilution), IBA1 (019-19741, FUJIFILM Wako Pure Chemical Corporation; 1:500 dilution), GFAP (AB5804, Abcam; 1:500 dilution), S100β (ab41548, Abcam; 1:300 dilution), CD68 (MCA1957, Bio-Rad Antibodies; 1:200 dilution), TDP-43 (12892-1-AP, Proteintech; 1:400 dilution), COL6 (ab182744, Abcam; 1:200), CTIP2 (ab18465, Abcam; 1:500).

Images were taken using a Zeiss LSM 880 confocal microscope or ZEISS Axio Scan.Z1 slide scanner. Image analyses were performed using ImageJ, Imaris, or QuPath-0.3.2 software.

### Surgical procedures for *in-vivo* recordings

Surgical and experimental procedures were conducted in accordance with the Animal (Scientific procedures) Act 1986, approved by the Animal Welfare and Ethical Review Body (AWERB) at University College London (UCL), and performed under an approved UK Home Office project licence at UCL. Mice were maintained in a 12 light/dark cycle with food and water supplied ad libitum. Prior to surgical procedures, WT, PR(400) and GR(400) mice were anaesthetised with isoflurane (3-4% induction, 1.5-2% for maintenance) and given a subcutaneous carprofen for pain relief during subsequent surgery. Liquid eye gel (Viscotears) was applied to protect the eyes during procedures and the animal’s head was shaved to remove fur. The animal was placed in a stereotaxic frame (WPI) and over a heating blanket to maintain normothermia during procedures. Exposed skin was disinfected using diluted chlorhexidine and cleaned with ethanol, following which lidocaine/prilocaine cream was gently applied to skin using a sterile applicator. All surgical procedures were performed under aseptic conditions. Anaesthetic depth was confirmed by monitoring the pedal reflex and breathing rate. Subcutaneous saline (0.2 ml) was provided every hour of surgery to maintain hydration. Following a small incision, the skin overlying the skull was carefully retracted and connective tissue over the skull carefully cleared. A small craniotomy overlying motor cortex was then performed using a hand-held microdrill (WPI) under constant cooling with sterile phosphate buffered saline and visualisation through a surgical microscope (Leica). In a subset of animals, a silver chloride wire was then attached to a small indentation in the skull overlying the cerebellum using cyanoacrylate glue (to act as a ground/reference electrode) and dental cement (Jet) used to secure the wire and build a well encircling the craniotomy. These animals were then transferred directly to the electrophysiology station while remaining in the stereotactic frame for subsequent acute recordings with no interruption of anaesthesia. In remaining animals, A glass coverslip (5 mm diameter) was placed over the craniotomy, and dental cement (Jet) applied to secure the cranial window and cover remaining exposed skull. On completion of these procedures, a subcutaneous injection of buprenorphine (0.1 mg/kg) was administered for immediate post-surgical pain relief, and the animal allowed to recover from surgical anaesthesia on a heated plate (37°C) and monitored continuously. On recovery, animals were returned to the holding room in single-housed conditions and carprofen provided in drinking water overnight and continued for three days, during which the animal’s health and welfare was closely monitored. Recordings were performed at least two weeks following recovery.

### In vivo two-photon Ca^2+^ imaging and analysis

Animals were initially anaesthetised with isoflurane (3-4% induction) and placed in a stereotaxic frame and over a heating blanket to maintain core temperature at 37°C throughout experimental procedures. In vivo two-photon Ca^2+^ imaging of motor cortex was performed under light isoflurane anesthesia (∼1%) using a custom-built resonant-scanning two-photon microscope (Independent NeuroScience Services) controlled by ScanImage (MBF Bioscience), and equipped with a Coherent Chameleon Discovery NX tunable laser and a 16×, 0.8 NA, Nikon water immersion objective. Imaging of neuronal calcium activity was performed at a wavelength of 1070 nm and fluorescence detected with a GaAsP photomultiplier tube (Hamamatsu). Images (512×512 pixels) were acquired at 30 Hz frame rate, and each field-of-view (FOV) was recorded for at least 5 minutes. Image analysis was performed with Suite2p ^92^ and custom MATLAB scripts. The recorded image stacks were first loaded into Suite2p for motion correction, region of interest (ROI) detection, and calcium signal extraction. For each detected ROI (putative cell somata), the neuropil corrected signal was extracted by subtracting the neuropil fluorescence signal surrounding the ROI (F_n_) from the raw fluorescence signal within the ROI (F): F_corr_(t)=F(t)-0.7*F_n_(t). The baseline fluorescence (F_0_) was estimated by using robust mean estimation and relative fluorescence change (ΔF/F=(F_corr_(t)-F_0_)/F_0_) over time was generated for each ROI. Ca^2+^ transients were identified as relative changes in ΔF/F that were two times larger than the standard deviation of the noise band. Following automatic peak detection, the peaks were inspected using custom-written MATLAB scripts and manually curated to exclude false positives and include false negatives. Silent and hyperactive neurons were defined as those with individual activity rates of 0 and >3 transients per min, respectively.

### *In vivo* Neuropixels recordings and analysis

For recordings following surgical recovery, animals were anaesthetised with isoflurane (3-4% induction) and placed in a stereotaxic frame and over a heating blanket to maintain core temperature at 37°C throughout experimental procedures. A small aperture overlying the motor cortex was made in the glass cover slip using a diamond-tipped drill bit and micro-drill, and a silver chloride ground/reference electrode affixed to the edge of the cranial window using dental cement (Jet). In all animals, a Neuropixels probe ^93^ (IMEC), was connected to the ground/reference electrode and slowly implanted into motor cortex using stereotactic procedures at a rate ∼5-10 µm/s under remote micromanipulator control (QUAD, Sutter Instruments) and visualised through a microscope (GT Vision). Following successful implantation, the cranial well was filled with warm sterile saline and the brain allowed to rest for at least 45 min before recordings, during which isoflurane anaesthesia was gradually lowered to maintain adequate anaesthetic depth and promote physiological cortical activity levels. A 10 minute recording of spontaneous brain activity was then performed in each animal. Following recordings, all animals were immediately sacrificed without recovery using a Schedule 1 method. Neuropixels recordings (30kHz sampling rate) of spontaneous brain activity were processed and automatically spike sorted using Kilosort3 using default parameters ^94^. Processed data were then imported into PHY software (https://github.com/cortex-lab/phy) for interactive visualisation of putative cortical clusters, and data manually inspected and curated to exclude false positives and include false negatives, and improve clustering through merging where appropriate. Following curation, spike-sorted data were analysed using custom-written MATLAB scripts and subjected to an additional quality control where only units in which less than 1% of associated spikes violated the physiological refractory period of 2 ms were included for further analysis. Mean firing rates of all units (as log_10_Hz) across the entire 10 minute recording session were calculated over 1 second time-bins for superficial cortex (0-400µm below surface) and deep layer 5 (550-800 µm below surface). Local field potential (LFP) data for the entire 10 minute recording session was low pass filtered and decimated from 2.5kHz to 500Hz using an 8th order Chebyshev Type 1 IIR filter and common average referenced. LFP data was averaged across cortical channels and mean power in the slow (0.1-1Hz), mid-gamma (40-90Hz) and high-gamma (90-120Hz) frequency bands estimated using Welch’s technique (40s windows with 50% overlap, MATLAB function ‘pwelch’). Data were tested for normal distributions using Kolmogorov-Smirnov or Anderson-Darling tests.

### Locomotor, grip strength and body weight assessment

Behavioural tests were performed monthly from 3-to 9-months of age, and in 12-month old mice. Motor coordination was measured by rotarod analysis (Ugo Basile). Mice were trained the week before starting the test. Then, mice received a session which included three trials of accelerated rotarod for a maximum of 300 seconds. Trials started at 4 rpm speed and accelerated up to 40 rpm in 4 min, the final min of the test was performed at 40 rpm. The average of recordings for each mouse was used to analyse rotarod performance. For grip strength analysis of muscle force, a grip strength meter (Bioseb) was used to measure forelimb and hindlimb grip strength. The grip strength meter was positioned, and the mice were held by the tail and lowered toward the apparatus. Mice were allowed to grasp the smooth metal grid with their forelimbs and hindlimbs and then were pulled backward. The force applied to the grid, measured at the moment in which the grasp was released, was recorded as the peak tension. The highest muscle force score of three independent trials was used. Body weight was measured weekly from 3-months of age. Mice were randomized into different experimental groups and the operator was blind to genotype.

### Motor neuron counts

10 μm-thick OCT-embedded spinal cord sections were stained with Cresyl Violet and motor neurons located within the sciatic motor pool were counted in each ventral horn on 35 sections, collected every 60 μm of tissue, covering L3 to L5 levels of the spinal cord.

### *In vivo* isometric muscle tension physiology

Isometric muscle tension physiology was performed as previously described ^95, 96^. Briefly, under deep anaesthesia (isoflurane inhalation via nose cone), hindlimbs were immobilized and the distal tendons of the TA and EDL muscles of both hindlimbs were exposed and consecutively attached to force transducers in parallel. Sciatic nerves were exposed bilaterally, at mid-thigh level, severed and the distal stumps placed in contact with stimulating electrodes. EDL muscle motor unit number estimates (MUNE) were determined by gradually increasing the amplitude of repeated square wave stimuli, thereby inducing stochastic changes in contractile force. The total number of motor units recruited over the full range of amplitudes was counted for individual muscles and averaged for each genotype.

### Proteomic analysis

Mouse lumbar spinal cords were solubilized in SDS-Lysis buffer (2% (wt/vol) SDS in 100 mM TEABC supplemented with Roche protease mini and Phos-STOP cocktail tablets). Automated homogenization was performed using the Precellys evolution homogenizer (Bertin Technologies # P000062-PEVO0-A). The lysates were then subjected to sonication using an automated Diagenode Bioruptor sonicator. Lysates were then centrifuged at 20800 g for 20 min and the supernatantused for. protein estimation by BCA assay and protein quality confirmed by SDS-PAGE.

200 µg of lysate was aliquoted and processed using S-Trap assisted On-column tryptic digestion as described previously (<u>dx.doi.org/10.17504/protocols.io.bs3tngnn</u>).. Peptides were eluted off the column by adding 80 µl of 50mM TEABC, 80 µl of 0.15% (vol/vol) formic acid in H2O and 3 X 80 µl of elution buffer containing 80% (vol/vol) in 0.15% (vol/vol) formic acid. The eluates were then subjected to vacuum drying using a Speedvac concentrator and stored at –80°C until further analysis. Following, TMT labelling and High-pH fractionation for LC-MS/MS analysis was carried out. A total of 96 fractions were collected and pooled to 48 fractions, vacuum dried and stored at –80°C until LC-MS/MS analysis.

LC-MS/MS analysis: A total of 48 bRPLC fractions were prepared for mass spectrometry analysis. 5% of the peptide digest was transferred into the autosampler vials and analysed using an Orbitrap Tribrid Lumos mass spectrometer with an Ultimate 3000 RSLC nano-liquid chromatography system. Samples were loaded on pre-column (C18, 5 µm, 100 A°, 100 µ, 2 cm Nano-viper column # 164564, Thermo Scientific) at 5 µl/min flow rate for about 10 minutes and then peptides were separated on a 50 cm column (C18, 5 µm, 50 cm, 100 A° Easy nano spray column # ES903, Thermo Scientific) by applying a non-linear gradient. LC was operated at 300 nl/min flow rate with a total run time of 100 min for each fraction. The mass spectrometer was operated in a data-dependent SPS-MS3 mode in a top speed for 2 seconds. Full Scan was acquired at 120K resolution at 200 m/z measured using Orbitrap mass analyser in the scan range of 350-1500 m/z. The data dependent MS2 scans were isolated using quadrupole mass filter with 0.7 Th isolation width and fragmented using normalised 32% HCD and detected using an Ion trap mass analyser which was operated in a rapid mode. Synchronous precursor selection of top 10 fragment ions further isolated and fragmented using normalised 55% HCD energy and measured using Orbitrap mass analyser at 55K resolution at 200 m/z in the scan range of 100-500. The AGC target and Ion injection times were set as 2e5, 50ms for MS1 and 10e4, 50ms for MS2 and 1e5, 120ms for MS3. Mono isotopic precursor selection (MIPS) was enabled, and charge state was set between 2 to 7 and dynamic exclusion duration was set at 45 sec.

Data analysis for mouse tissue: Spinal cord raw MS data was searched with MaxQuant software suite (version: 2.0.1.0) ^97^ against the Uniprot Mouse database appended with (GR)400 and (PR)400 sequences for C9orf72 and a common contaminant list exists within MaxQuant. The following parameters were used: Carbamidomethylation of Cys as fixed modification; Oxidation of Met, deamidation of Asn and Gln; Acetylation of Protein-N-ter and Phosphorylation of Ser/Thr/Tyr were set as variable modifications. Trypsin was set as a protease with a maximum of two missed cleavages allowed and a minimum peptide length of 7 amino acids. FDR was set at 1% for both protein and PSM level. The protein group output files were further processed using Perseus software suite (version: 1.6.15.0) ^98^ for downstream statistical analysis. Welch’s t-Test was carried out between the groups to identify differentially regulated proteins. Gene-Ontology analysis was performed on differentially regulated proteins using enrichR software ^99^.

Reanalysis of C9orf72 iPSC-derived motor neurons: we downloaded the SWATH MS raw data from the CHORUS repository containing 10 control and 7 ALS samples. The .wiff and .wiff.scan files were then converted to mzML using MS MSconvert with the peak picking filter added ^100^.Converted files were then searched using Spectronaut software suite Version Rubin: 15.7.220308.50606 ^101^ using a direct-DIA strategy. A FASTA file from the Human UniProt database was used to generate a predicted library within Spectronaut and search was performed using Pulsar search algorithm with a default search parameters. The output protein group file was then processed using Perseus software suite as described above to perform Welch’s t-Test to identify differentially regulated proteins between ALS and control samples. GO analysis was performed using the enrichR software suite. The search was performed using the default settings within DIA-NN.

### Microarray analysis

We analysed the transcriptional signature in laser-captured spinal motor neurons from postmortem C9-ALS patients ^51^. Raw microarray data is available in Gene Expression Omnibus (GEO) with accession number GSE56504. We performed Gene Set Enrichment Analysis (GSEA) using the gseGO function from the clusterProfiler R package ^102^. Differentially expressed genes were ranked and then subjected to GSEA. The top enriched gene sets included all had normalized enrichment score (NES) > 0).

### Ingenuity Pathway Analysis

Data were analysed with the use of QIAGEN IPA (QIAGEN Inc., https://digitalinsights.qiagen.com/IPA). We used userdata set as the reference data set and p=0.05 and LFC –1.5-1.5.

### Human frontal cortex RNAseq data analysis

We used previously published RNAseq data from the frontal cortex of pathologically diagnosed FTLD patients (with and without MND) and control samples. We extracted the summary statistics and residual expression values for our gene(s) or pathways of interest from the differential gene expression and Weighted Gene Co-expression Network Analysis (WGCNA) with adjustment for cell-type markers ^52^.

### iPSC differentiation and lentiviral transduction

The WTC11 iPSC line were differentiated into cortical neurons (i^3^Neurons) using a method previously described ^54^. Briefly, iPSCs were grown to 70-80% confluency. On DIV 0 of neuronal induction, cells were washed with PBS and singularized with Accutase (Gibco). Cells were plated at a desired ratio (typically 3.75×10^5^ cells/well of a 6-well plate) onto Geltrex-coated plates. Cells were maintained in DMEM-F12 (Gibco) containing 1× N2 (Thermo Fisher Scientific), 1× Glutamine (Gibco), 1× HEPES (Gibco), 1× NEAA (Gibco), doxycycline (2µg/mL) and 10 µM Y-27632 (day 0 only; Tocris). Media was changed every day for 3 days. On DIV 3, neural progenitor cells were dissociated with accutase and replated onto poly-L-ornithine (Merck) and laminin-coated plates in neuronal maintenance media: Neurobasal (Gibco), supplemented with 1× B27 (Gibco), 10 ng/mL BDNF (PeproTech), 10 ng/mL NT-3 (PreproTech) and 1 µg/mL laminin. Neurons were plated at a desired ratio (typically 6×10^5^ cells/well of a 6-well plate).On DIV 3, lentivirus was added to i^3^Neurons to overexpress C9orf72(GR)_50_ or GFP constructs under the control of the neuron-specific human synapsin 1 promoter (hSYN) as previously described ^55^. From DIV 3 to DIV 7, cells were maintained in neuronal maintenance media and imaged on the Incucyte^®^ S3 Live-Cell Analysis System using a confluency mask to quantify and track cell survival.

### Drosophila stocks and maintenance

*Drosophila* stocks were maintained on SYA food (15 g/L agar, 50 g/L sugar, 100 g/L brewer’s yeast, 30 ml/L nipagin [10% in ethanol], and 3 ml/L propionic acid) at 25°C in a 12 h light/dark cycle with 60% humidity. The UAS-(GR)36 flies have been previously described ^15^. The fly lines GMR-Gal4 (Bloomington #9146), UAS-*daw*^RNAi^ (Bloomington #50911), and UAS-Mp^RNAi^ (Bloomington #52981) were obtained from the Bloomington Drosophila Stock Centre.

*Drosophila* ortholog prediction: The Drosophila RNAi Screening Center Integrative Ortholog Prediction Tool (DIOPT; http://www.flyrnai.org/diopt) was used to search for orthologues of TGF-β1, COL6A1, COL6A2, and COL6A3. DIOPT predicted *daw* as the *Drosophila* orthologue of *TGF-β1*, and Mp as the *Drosophila* orthologue of human COL6A1, COL6A2 and COL6A3.

Assessment of eye phenotypes: Flies carrying the UAS-*daw*^RNAi^ or UAS-*Mp*^RNAi^ construct were crossed to the GMR-GAL4; GMR-GR36 driver line at 25°C. Two-day old adult progeny were imaged using a stereomicroscope, with female eyes used for analyses. All eye images were obtained under the same magnification; eye area was calculated from each image using ImageJ ^103^.

### Statistical analysis

All data are presented as mean ± standard error of the mean (SEM). Statistical differences of continuous data from two experimental groups were calculated using unpaired two-sample Student’s t-test. Comparisons of data from more than two groups were performed using a one-way-ANOVA followed by Bonferroni correction for multiple comparisons. When two independent variables were available, comparisons of data from more than two groups were performed using a two-way-ANOVA followed by Bonferroni correction for multiple comparisons. Statistical significance threshold was set at P < 0.05, unless otherwise indicated. Statistical methods were used to predetermine sample sizes.

### Data collection

Data collection and analysis were performed blind to the conditions of the experiments. Data available upon request.

## Acknowledgements

We thank Manuela Neumann for providing the C9orf72 antibody and Aaron Gitler and Maya Maor-Nof for sharing the GR_50_ lentiviral construct. We also thank the UK DRI at UCL animal technicians Elena Ghirardello and Phill Muckett, and the Biological Services Units at UCL, for assisting with the care and welfare of our animals.

## References

1. Hardiman, O. et al. Amyotrophic lateral sclerosis. Nat Rev Dis Primers 3, 17085 (2017).

2. Olney, N. T., Spina, S. & Miller, B. L. Frontotemporal Dementia. Neurol Clin 35, 339–374 (2017).

3. DeJesus-Hernandez, M. et al. Expanded GGGGCC hexanucleotide repeat in noncoding region of C9ORF72 causes chromosome 9p-linked FTD and ALS. Neuron 72, 245–256 (2011).

4. Renton, A. E. et al. A hexanucleotide repeat expansion in C9ORF72 is the cause of chromosome 9p21-linked ALS-FTD. Neuron 72, 257–268 (2011).

5. Majounie, E. et al. Frequency of the C9orf72 hexanucleotide repeat expansion in patients with amyotrophic lateral sclerosis and frontotemporal dementia: a cross-sectional study. Lancet Neurol 11, 323–330 (2012).

6. Rohrer, J. D. et al. C9orf72 expansions in frontotemporal dementia and amyotrophic lateral sclerosis. Lancet Neurol 14, 291–301 (2015).

7. Zu, T. et al. Non-ATG-initiated translation directed by microsatellite expansions. Proc Natl Acad Sci U S A 108, 260–265 (2011).

8. MacKenzie, I. R. et al. Dipeptide repeat protein pathology in C9ORF72 mutation cases: clinico-pathological correlations. Acta Neuropathol 126, 859–879 (2013).

9. Ash, P. E. A. et al. Unconventional translation of C9ORF72 GGGGCC expansion generates insoluble polypeptides specific to c9FTD/ALS. Neuron 77, 639–646 (2013).

10. Gendron, T. F. et al. Antisense transcripts of the expanded C9ORF72 hexanucleotide repeat form nuclear RNA foci and undergo repeat-associated non-ATG translation in c9FTD/ALS. Acta Neuropathol 126, 829–844 (2013).

11. Mori, K. et al. Bidirectional transcripts of the expanded C9orf72 hexanucleotide repeat are translated into aggregating dipeptide repeat proteins. Acta Neuropathol 126, 881–893 (2013).

12. Mori, K. et al. The C9orf72 GGGGCC repeat is translated into aggregating dipeptide-repeat proteins in FTLD/ALS. Science 339, 1335–1338 (2013).

13. Zu, T. et al. RAN proteins and RNA foci from antisense transcripts in C9ORF72 ALS and frontotemporal dementia. Proc Natl Acad Sci U S A 110, E4968–77 (2013).

14. Balendra, R. & Isaacs, A. M. C9orf72-mediated ALS and FTD: multiple pathways to disease. Nat Rev Neurol 14, 544–558 (2018).

15. Mizielinska, S. et al. C9orf72 repeat expansions cause neurodegeneration in Drosophila through arginine-rich proteins. Science 345, 1192–1194 (2014).

16. Wen, X. et al. Antisense proline-arginine RAN dipeptides linked to C9ORF72-ALS/FTD form toxic nuclear aggregates that initiate in vitro and in vivo neuronal death. Neuron 84, 1213– 1225 (2014).

17. Yang, D. et al. FTD/ALS-associated poly(GR) protein impairs the Notch pathway and is recruited by poly(GA) into cytoplasmic inclusions. Acta Neuropathol 130, 525–535 (2015).

18. Kwon, I. et al. Poly-dipeptides encoded by the C9orf72 repeats bind nucleoli, impede RNA biogenesis, and kill cells. Science 345, 1139–1145 (2014).

19. Tao, Z. et al. Nucleolar stress and impaired stress granule formation contribute to C9orf72 RAN translation-induced cytotoxicity. Hum Mol Genet 24, 2426–2441 (2015).

20. Lopez-Gonzalez, R. et al. Poly(GR) in C9ORF72-Related ALS/FTD Compromises Mitochondrial Function and Increases Oxidative Stress and DNA Damage in iPSC-Derived Motor Neurons. Neuron 92, 383–391 (2016).

21. Farg, M. A., Konopka, A., Soo, K. Y., Ito, D. & Atkin, J. D. The DNA damage response (DDR) is induced by the C9orf72 repeat expansion in amyotrophic lateral sclerosis. Hum Mol Genet 26, 2882–2896 (2017).

22. Verdone, B. M. et al. A mouse model with widespread expression of the C9orf72-linked glycine-arginine dipeptide displays non-lethal ALS/FTD-like phenotypes. Sci Rep 12, 5644 (2022).

23. Zhang, Y. J. et al. Poly(GR) impairs protein translation and stress granule dynamics in C9orf72-associated frontotemporal dementia and amyotrophic lateral sclerosis. Nat Med 24, 1136– 1142 (2018).

24. Cook, C. N. et al. C9orf72 poly(GR) aggregation induces TDP-43 proteinopathy. Sci Transl Med 12, eabb3774 (2020).

25. Choi, S. Y. et al. C9ORF72-ALS/FTD-associated poly(GR) binds Atp5a1 and compromises mitochondrial function in vivo. Nat Neurosci 22, 851–862 (2019).

26. Zhang, Y. J. et al. Heterochromatin anomalies and double-stranded RNA accumulation underlie C9orf72 poly(PR) toxicity. Science 363, eaav2606 (2019).

27. Hao, Z. et al. Motor dysfunction and neurodegeneration in a C9orf72 mouse line expressing poly-PR. Nat Commun 10, 2906 (2019).

28. LaClair, K. D. et al. Congenic expression of poly-GA but not poly-PR in mice triggers selective neuron loss and interferon responses found in C9orf72 ALS. Acta Neuropathol 140, 121–142 (2020).

29. Moens, T. G., Partridge, L. & Isaacs, A. M. Genetic models of C9orf72: what is toxic? Curr Opin Genet Dev 44, 92–101 (2017).

30. Todd, T. W. & Petrucelli, L. Modelling amyotrophic lateral sclerosis in rodents. Nature Reviews Neuroscience 2022, 1–21 (2022) doi:10.1038/s41583-022-00564-x.

31. O’Rourke, J. G. et al. C9orf72 is required for proper macrophage and microglial function in mice. Science 351, 1324–1329 (2016).

32. Atanasio, A. et al. C9orf72 ablation causes immune dysregulation characterized by leukocyte expansion, autoantibody production, and glomerulonephropathy in mice. Sci Rep 6, (2016).

33. Jiang, J. et al. Gain of Toxicity from ALS/FTD-Linked Repeat Expansions in C9ORF72 Is Alleviated by Antisense Oligonucleotides Targeting GGGGCC-Containing RNAs. Neuron 90, 535–550 (2016).

34. Burberry, A. et al. Loss-of-function mutations in the C9ORF72 mouse ortholog cause fatal autoimmune disease. Sci Transl Med 8, 347ra93 (2016).

35. Burberry, A. et al. C9orf72 suppresses systemic and neural inflammation induced by gut bacteria. Nature 582, 89–94 (2020).

36. Koppers, M. et al. C9orf72 ablation in mice does not cause motor neuron degeneration or motor deficits. Ann Neurol 78, 426–438 (2015).

37. Sullivan, P. M. et al. The ALS/FTLD associated protein C9orf72 associates with SMCR8 and WDR41 to regulate the autophagy-lysosome pathway. Acta Neuropathol Commun 4, 51 (2016).

38. Sudria-Lopez, E. et al. Full ablation of C9orf72 in mice causes immune system-related pathology and neoplastic events but no motor neuron defects. Acta Neuropathol 132, 145– 147 (2016).

39. Chew, J. et al. Neurodegeneration. C9ORF72 repeat expansions in mice cause TDP-43 pathology, neuronal loss, and behavioral deficits. Science 348, 1151–1154 (2015).

40. Chew, J. et al. Aberrant deposition of stress granule-resident proteins linked to C9orf72-associated TDP-43 proteinopathy. Mol Neurodegener 14, 9 (2019).

41. O’Rourke, J. G. et al. C9orf72 BAC Transgenic Mice Display Typical Pathologic Features of ALS/FTD. Neuron 88, 892–901 (2015).

42. Peters, O. M. et al. Human C9ORF72 Hexanucleotide Expansion Reproduces RNA Foci and Dipeptide Repeat Proteins but Not Neurodegeneration in BAC Transgenic Mice. Neuron 88, 902–909 (2015).

43. Liu, Y. et al. C9orf72 BAC Mouse Model with Motor Deficits and Neurodegenerative Features of ALS/FTD. Neuron 90, 521–534 (2016).

44. de Giorgio, F., Maduro, C., Fisher, E. M. C. & Acevedo-Arozena, A. Transgenic and physiological mouse models give insights into different aspects of amyotrophic lateral sclerosis. DMM Disease Models and Mechanisms 12, dmm037424 (2019).

45. Gao, F., Almeida, S. & Lopez-Gonzalez, R. Dysregulated molecular pathways in amyotrophic lateral sclerosis-frontotemporal dementia spectrum disorder. EMBO J 36, 2931–2950 (2017).

46. Ghaffari, L. T., Trotti, D., Haeusler, A. R. & Jensen, B. K. Breakdown of the central synapses in C9orf72-linked ALS/FTD. Front Mol Neurosci 15, 1005112 (2022).

47. Coyne, A. N. et al. G4C2 Repeat RNA Initiates a POM121-Mediated Reduction in Specific Nucleoporins in C9orf72 ALS/FTD. Neuron 107, 1124–1140.e11 (2020).

48. Meyer, D. E. & Chilkoti, A. Genetically encoded synthesis of protein-based polymers with precisely specified molecular weight and sequence by recursive directional ligation: examples from the elastin-like polypeptide system. Biomacromolecules 3, 357–367 (2002).

49. Phatnani, H. et al. An integrated multi-omic analysis of iPSC-derived motor neurons from C9ORF72 ALS patients. iScience 24, 103221 (2021).

50. Wong, C. O. & Venkatachalam, K. Motor neurons from ALS patients with mutations in C9ORF72 and SOD1 exhibit distinct transcriptional landscapes. Hum Mol Genet 28, 2799– 2810 (2019).

51. Highley, J. R. et al. Loss of nuclear TDP-43 in amyotrophic lateral sclerosis (ALS) causes altered expression of splicing machinery and widespread dysregulation of RNA splicing in motor neurones. Neuropathol Appl Neurobiol 40, 670–685 (2014).

52. Dickson, D. W. et al. Extensive transcriptomic study emphasizes importance of vesicular transport in C9orf72 expansion carriers. Acta Neuropathol Commun 7, 150 (2019).

53. Meng, X. M., Nikolic-Paterson, D. J. & Lan, H. Y. TGF-β: the master regulator of fibrosis. Nat Rev Nephrol 12, 325–338 (2016).

54. Fernandopulle, M. S. et al. Transcription-factor mediated differentiation of human iPSCs into neurons. Curr Protoc Cell Biol 79, e51 (2018).

55. Maor-Nof, M. et al. p53 is a central regulator driving neurodegeneration caused by C9orf72 poly(PR). Cell 184, 689–708.e20 (2021).

56. Atilano, M. L. et al. Enhanced insulin signalling ameliorates C9orf72 hexanucleotide repeat expansion toxicity in Drosophila. Elife 10, e58565 (2021).

57. Gendron, T. F. & Petrucelli, L. Disease Mechanisms of C9ORF72 Repeat Expansions. Cold Spring Harb Perspect Med 8, a024224 (2018).

58. Pasniceanu, I. S., Atwal, M. S., Souza, C. D. S., Ferraiuolo, L. & Livesey, M. R. Emerging Mechanisms Underpinning Neurophysiological Impairments in C9ORF72 Repeat Expansion-Mediated Amyotrophic Lateral Sclerosis/Frontotemporal Dementia. Front Cell Neurosci 15, 784833 (2021).

59. Sakae, N. et al. Poly-GR dipeptide repeat polymers correlate with neurodegeneration and Clinicopathological subtypes in C9ORF72-related brain disease. Acta Neuropathol Commun 6, 63 (2018).

60. Saberi, S. et al. Sense-encoded poly-GR dipeptide repeat proteins correlate to neurodegeneration and uniquely co-localize with TDP-43 in dendrites of repeat-expanded C9orf72 amyotrophic lateral sclerosis. Acta Neuropathol 135, 459–474 (2018).

61. Quaegebeur, A., Glaria, I., Lashley, T. & Isaacs, A. M. Soluble and insoluble dipeptide repeat protein measurements in C9orf72-frontotemporal dementia brains show regional differential solubility and correlation of poly-GR with clinical severity. Acta Neuropathol Commun 8, 184 (2020).

62. Jo, Y., Lee, J., Lee, S. Y., Kwon, I. & Cho, H. Poly-dipeptides produced from C9orf72 hexanucleotide repeats cause selective motor neuron hyperexcitability in ALS. Proc Natl Acad Sci U S A 119, e2113813119 (2022).

63. Suzuki, N. et al. The mouse C9ORF72 ortholog is enriched in neurons known to degenerate in ALS and FTD. Nat Neurosci 16, 1725–1727 (2013).

64. Langseth, A. J. et al. Cell-type specific differences in promoter activity of the ALS-linked C9orf72 mouse ortholog. Sci Rep 7, 5685 (2017).

65. Frick, P. et al. Novel antibodies reveal presynaptic localization of C9orf72 protein and reduced protein levels in C9orf72 mutation carriers. Acta Neuropathol Commun 6, 72 (2018).

66. Mizielinska, S. et al. C9orf72 frontotemporal lobar degeneration is characterised by frequent neuronal sense and antisense RNA foci. Acta Neuropathol 126, 845–857 (2013).

67. DeJesus-Hernandez, M. et al. In-depth clinico-pathological examination of RNA foci in a large cohort of C9ORF72 expansion carriers. Acta Neuropathol 134, 255–269 (2017).

68. Lall, D. et al. C9orf72 deficiency promotes microglial-mediated synaptic loss in aging and amyloid accumulation. Neuron 109, 2275–2291.e8 (2021).

69. Ziff, O. J. et al. Meta-analysis of human and mouse ALS astrocytes reveals multi-omic signatures of inflammatory reactive states. Genome Res 32, 71–84 (2022).

70. Humphrey, J. et al. Integrative transcriptomic analysis of the amyotrophic lateral sclerosis spinal cord implicates glial activation and suggests new risk genes. Nat Neurosci 26, 150–162 (2023).

71. Prehn, J. H. M. & Krieglstein, J. Opposing effects of transforming growth factor-beta 1 on glutamate neurotoxicity. Neuroscience 60, 7–10 (1994).

72. Ruocco, A. et al. A transforming growth factor-beta antagonist unmasks the neuroprotective role of this endogenous cytokine in excitotoxic and ischemic brain injury. J Cereb Blood Flow Metab 19, 1345–1353 (1999).

73. Boche, D., Cunningham, C., Gauldie, J. & Perry, V. H. Transforming growth factor-beta 1-mediated neuroprotection against excitotoxic injury in vivo. J Cereb Blood Flow Metab 23, 1174–1182 (2003).

74. Prehn, J. H. M., Backhauß, C. & Krieglstein, J. Transforming growth factor-beta 1 prevents glutamate neurotoxicity in rat neocortical cultures and protects mouse neocortex from ischemic injury in vivo. J Cereb Blood Flow Metab 13, 521–525 (1993).

75. McNeill, H. et al. Neuronal rescue with transforming growth factor-beta 1 after hypoxic-ischaemic brain injury. Neuroreport 5, 901–904 (1994).

76. Zhu, Y. et al. Transforming growth factor-beta 1 increases bad phosphorylation and protects neurons against damage. J Neurosci 22, 3898–3909 (2002).

77. Brionne, T. C., Tesseur, I., Masliah, E. & Wyss-Coray, T. Loss of TGF-β1 leads to increased neuronal cell death and microgliosis in mouse brain. Neuron 40, 1133–1145 (2003).

78. Koeglsperger, T. et al. Impaired glutamate recycling and GluN2B-mediated neuronal calcium overload in mice lacking TGF-β1 in the CNS. Glia 61, 985–1002 (2013).

79. Cheng, J. S. et al. Collagen VI protects neurons against Abeta toxicity. Nat Neurosci 12, 119– 121 (2009).

80. Cheng, I. H. et al. Collagen VI protects against neuronal apoptosis elicited by ultraviolet irradiation via an Akt/phosphatidylinositol 3-kinase signaling pathway. Neuroscience 183, 178–188 (2011).

81. Cescon, M., Gattazzo, F., Chen, P. & Bonaldo, P. Collagen VI at a glance. J Cell Sci 128, 3525– 3531 (2015).

82. Cescon, M., Chen, P., Castagnaro, S., Gregorio, I. & Bonaldo, P. Lack of collagen VI promotes neurodegeneration by impairing autophagy and inducing apoptosis during aging. Aging 8, 1083–1101 (2016).

83. Phatnani, H. P. et al. Intricate interplay between astrocytes and motor neurons in ALS. Proc Natl Acad Sci U S A 110, E756–65 (2013).

84. Meroni, M. et al. Transforming growth factor beta 1 signaling is altered in the spinal cord and muscle of amyotrophic lateral sclerosis mice and patients. Neurobiol Aging 82, 48–59 (2019).

85. Zubiri, I. et al. Tissue-enhanced plasma proteomic analysis for disease stratification in amyotrophic lateral sclerosis. Mol Neurodegener 13, 60 (2018).

86. Houi, K., Kobayashi, T., Kato, S., Mochio, S. & Inoue, K. Increased plasma TGF-beta1 in patients with amyotrophic lateral sclerosis. Acta Neurol Scand 106, 299–301 (2002).

87. Iłz̈ecka, J., Stelmasiak, Z. & Dobosz, B. Transforming growth factor-beta 1 (TGF-beta 1) in patients with amyotrophic lateral sclerosis. Cytokine 20, 239–243 (2002).

88. Masuda, T. et al. Transforming growth factor-β1 in the cerebrospinal fluid of patients with distinct neurodegenerative diseases. J Clin Neurosci 35, 47–49 (2017).

89. Galbiati, M. et al. Multiple Roles of Transforming Growth Factor Beta in Amyotrophic Lateral Sclerosis. Int J Mol Sci 21, 1–17 (2020).

90. Simone, R. et al. G-quadruplex-binding small molecules ameliorate C9orf72 FTD/ALS pathology in vitro and in vivo. EMBO Mol Med 10, 22–31 (2018).

91. Moens, T. G. et al. Sense and antisense RNA are not toxic in Drosophila models of C9orf72-associated ALS/FTD. Acta Neuropathol 135, 445–457 (2018).

92. Pachitariu, M., et al. Suite2p: beyond 10,000 neurons with standard two-photon microscopy. bioRxiv 61507 (2017) doi:10.1101/061507.

93. Jun, J. J. et al. Fully integrated silicon probes for high-density recording of neural activity. Nature 551, 232–236 (2017).

94. Pachitariu, M., Sridhar, S. & Stringer, C. Solving the spike sorting problem with Kilosort. bioRxiv 2023.01.07.523036 (2023) doi:10.1101/2023.01.07.523036.

95. Ahmed, M. et al. Targeting protein homeostasis in sporadic inclusion body myositis. Sci Transl Med 8, 331ra41 (2016).

96. Gray, A. L. et al. Deterioration of muscle force and contractile characteristics are early pathological events in spinal and bulbar muscular atrophy mice. Dis Model Mech 13, dmm042424 (2020).

97. Cox, J. & Mann, M. MaxQuant enables high peptide identification rates, individualized p.p.b.-range mass accuracies and proteome-wide protein quantification. Nat Biotechnol 26, 1367– 1372 (2008).

98. Tyanova, S. et al. The Perseus computational platform for comprehensive analysis of (prote)omics data. Nat Methods 13, 731–740 (2016).

99. Kuleshov, M. v., et al. Enrichr: a comprehensive gene set enrichment analysis web server 2016 update. Nucleic Acids Res 44, W90–W97 (2016).

100. Chambers, M. C. et al. A cross-platform toolkit for mass spectrometry and proteomics. Nat Biotechnol 30, 918–920 (2012).

101. Bruderer, R. et al. Extending the limits of quantitative proteome profiling with data-independent acquisition and application to acetaminophen-treated three-dimensional liver microtissues. Mol Cell Proteomics 14, 1400–1410 (2015).

102. Yu, G., Wang, L. G., Han, Y. & He, Q. Y. ClusterProfiler: An R package for comparing biological themes among gene clusters. OMICS 16, 284–287 (2012).

103. Schneider, C. A., Rasband, W. S. & Eliceiri, K. W. NIH Image to ImageJ: 25 years of Image Analysis. Nat Methods 9, 671 (2012).

