## Supplemental Figures for "PolyGR and polyPR knock-in mice reveal a conserved neuroprotective extracellular matrix signature in *C9orf72* ALS/FTD neurons"

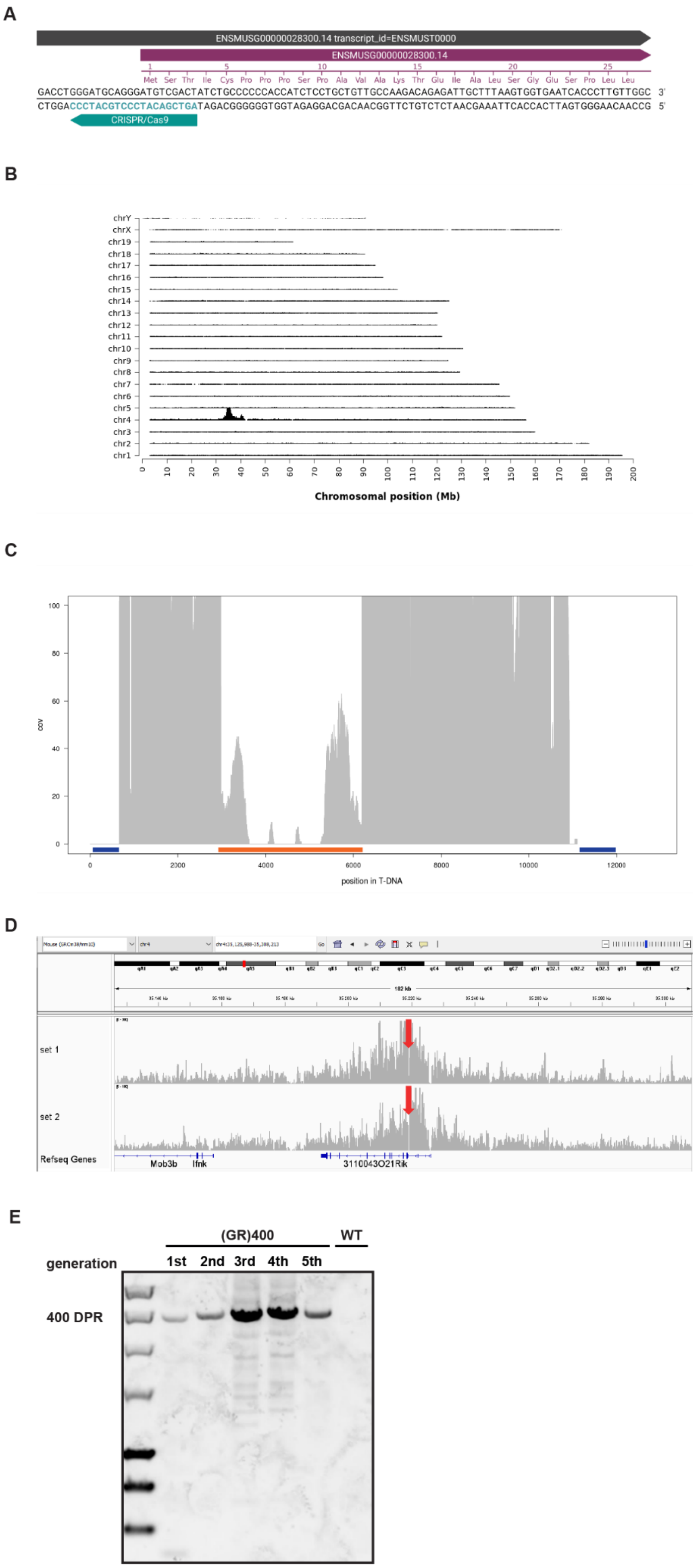
A
ENSMUSG00000028300.14 transcript_id=ENSMUST0000
ENSMUSG00000028300.14
Met Ser Thr Ile Cys Pro Pro Pro Ser Pro Ala Val Ala Lys Thr Glu Ile Ala Leu Ser Gly Glu Ser Pro Leu Leu
CRISPR/Cas9
B
Chromosomal position (Mb)
C
cov
position in T-DNA
D
set 1
set 2
Refseq Genes
Mob3b
Ifnk
3110043O21Rik
E
(GR)400
WT
generation
1st
2nd
3rd
4th
5th
400 DPR

Supplementary Figure 1. **C9orf72 knock-in mice targeting strategy and confirmaton**

(A) Design for CRISPR assisted *C9orf72* gene targeting. The sgRNA for CRISPR/Cas9 is indicated by the teal bar.

(B) Targeted locus amplification sequence coverage across the mouse genome. The chromosomes are indicated on the y-axis, the chromosomal position on the x-axis.

(C) Targeted locus amplification sequence coverage across the knock-in sequence. The whole knock-in sequence has good coverage except for the blue underlined backbone sequences as expected from a correct targeting event. A coverage gap is present at knock-in:3522-5927, representative the 400 DPRs, and is underlined by the orange bar.

(D) Targeted locus amplification sequence coverage across the knock-in integration locus. The red arrow points toward the knock-in integration site.

(E) Repeat-length PCR of (GR)400 genomic DNA to establish repeat length and stability over five generation.

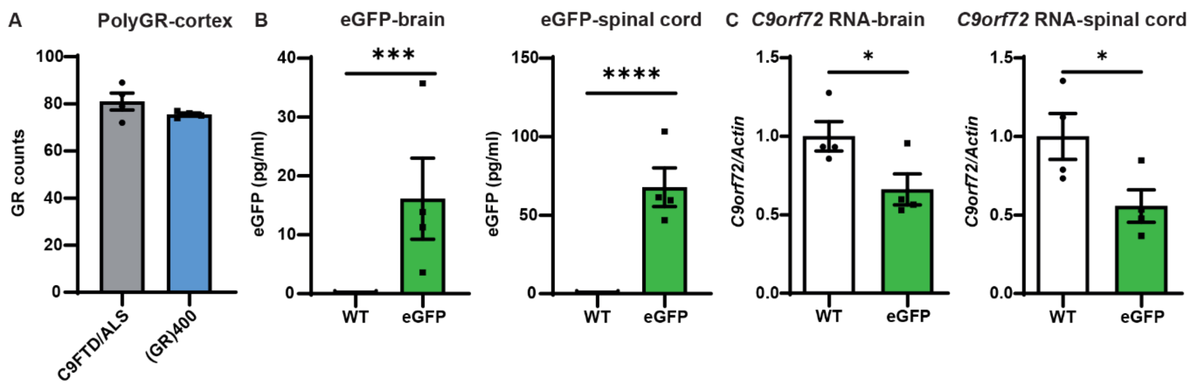


Supplementary Figure 2. **Patient and knock-in cortex polyGR comparison and generation of control *C9orf72*-eGFP knock-in mice**

(A) Quantification of polyGR proteins in C9FTD/ALS patient and 12-month old (GR)400 mice cortex by MSD immunoassay. Graph, mean ± SEM, n = 4 independent samples per experimental group, unpaired two-sample Student's t-test.

(B) Quantification of eGFP proteins in brain (left panel) and spinal cord (right panel) of WT and eGFP mice at 3 months of age by ELISA. Graph, mean ± SEM, n = 4 mice per genotype, unpaired two-sample Student's t-test, ***$p < 0.001$, ****$p < 0.0001$.

(C) qPCR analysis of *C9orf72* transcript levels normalised to *β-actin* in brain (left panel) and spinal cord (right panel) of WT and eGFP mice at 3 months of age. Graph, mean ± SEM, n = 4 mice per genotype, unpaired two-sample Student's t-test, *$p < 0.05$.

13

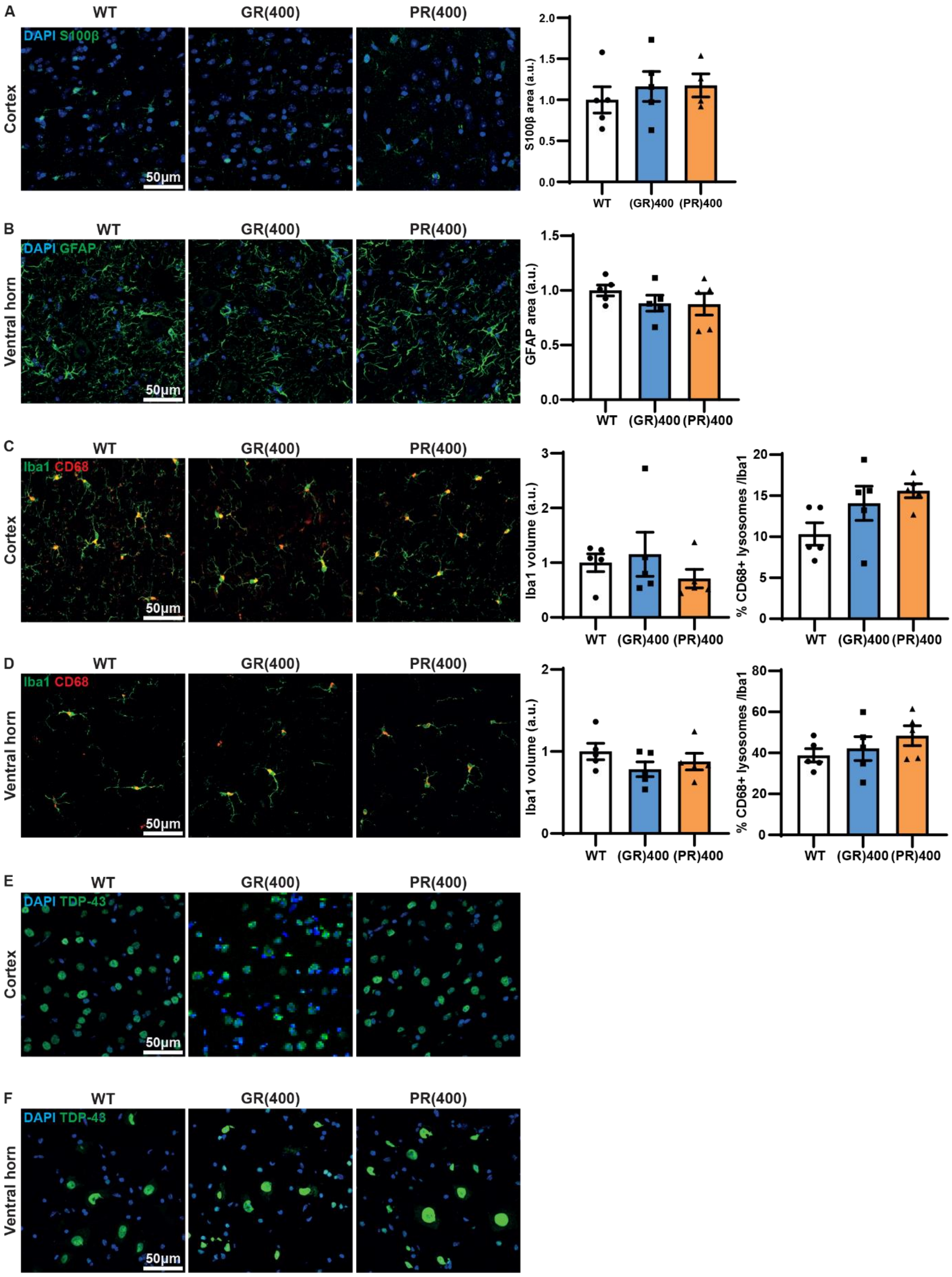

A
WT
GR(400)
PR(400)
DAPI S100β
Cortex
50µm
S100β area (a.u.)
2.0
1.5
1.0
0.5
0.0
WT (GR)400 (PR)400
B
WT
GR(400)
PR(400)
DAPI GFAP
Ventral horn
50µm
GFAP area (a.u.)
1.5
1.0
0.5
0.0
WT (GR)400 (PR)400
C
WT
GR(400)
PR(400)
Iba1 CD68
Cortex
50µm
Iba1 volume (a.u.)
3
2
1
0
WT (GR)400 (PR)400
% CD68+ lysosomes /Iba1
20
15
10
5
0
WT (GR)400 (PR)400
D
WT
GR(400)
PR(400)
Iba1 CD68
Ventral horn
50µm
Iba1 volume (a.u.)
2
1
0
WT (GR)400 (PR)400
% CD68+ lysosomes /Iba1
80
60
40
20
0
WT (GR)400 (PR)400
E
WT
GR(400)
PR(400)
DAPI TDP-43
Cortex
50µm
F
WT
GR(400)
PR(400)
DAPI TDP-43
Ventral horn
50µm

Supplementary Figure 3. **(GR)400 and (PR)400 knock-in mice do not exhibit gliosis in brain and spinal cord up to 12 months of age**

(A) Representative confocal images and quantification of immunofluorescence staining of astrocytic marker S100β (green) in brain cortex in WT, (GR)400, and (PR)400 mice at 12 months of age. DAPI (blue) stains nuclei. Graph, mean ± SEM, n = 5 mice per genotype, one-way ANOVA, Bonferroni's multiple comparison.

(B) Representative confocal images and quantification of immunofluorescence staining of astrocytic marker GFAP (green) in lumbar spinal cord ventral horn in WT, (GR)400, and (PR)400 mice at 12 months of age. DAPI (blue) stains nuclei. Graph, mean ± SEM, n = 5 mice per genotype, one-way ANOVA, Bonferroni's multiple comparison.

(C) Representative confocal images and quantification of immunofluorescence staining showing microglial density and colocalization between microglial markers Iba1 (green) and microglial lysosomal marker CD68 (red) in brain cortex in WT, (GR)400, and (PR)400 mice at 12 months of age. Graph, mean ± SEM, n = 5 mice per genotype, one-way ANOVA, Bonferroni's multiple comparison.

(D) Representative confocal images and quantification of immunofluorescence staining showing microglial density and colocalization between microglial markers Iba1 (green) and microglial lysosomal marker CD68 (red) in lumbar spinal cord ventral horn in WT, (GR)400, and (PR)400 mice at 12 months of age. Graph, mean ± SEM, n = 5 mice per genotype, one-way ANOVA, Bonferroni's multiple comparison.

(E) Representative confocal images of immunofluorescence staining showing TDP-43 (green) cellular localisation in brain cortex in WT, (GR)400, and (PR)400 mice at 12 months of age. DAPI (blue) stains nuclei.

(F) Representative confocal images of immunofluorescence staining showing TDP-43 (green) cellular localisation in lumbar spinal cord ventral horn in WT, (GR)400, and (PR)400 mice at 12 months of age. DAPI (blue) stains nuclei.

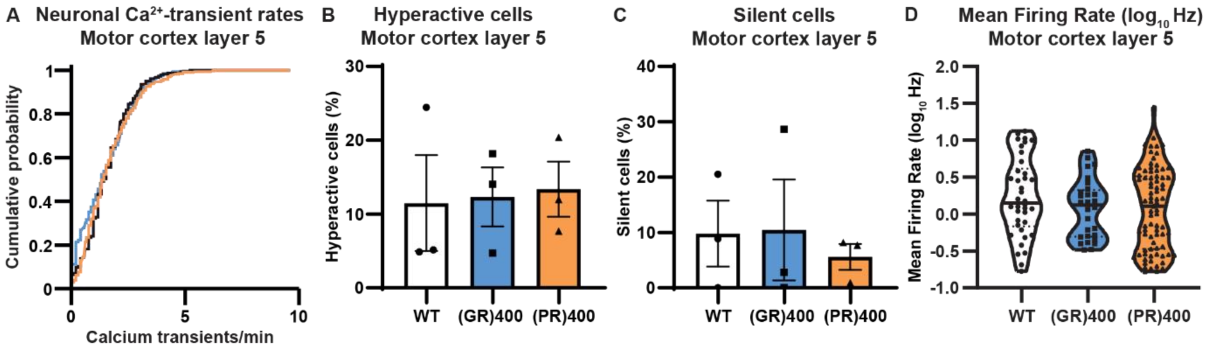


Supplementary Figure 4. **(GR)400 and (PR)400 knock-in mice do not exhibit cortical hyperexcitability in motor cortex layer 5**

(A) Cumulative distribution plot displaying neuronal $Ca^{2+}$-transient rates across animals in layer 5 of WT (187 cells, 4 mice), (GR)400 (616 cells, 3 mice) and (PR)400 (305 cells, 4 mice) mice at 15-19 months of age.

(B) Percentage of hyperactive (>3 $Ca^{2+}$-transients per minute) neurons in motor cortex layer 5 of WT, (GR)400, and (PR)400 mice at 15-19 months of age. Graph, mean ± SEM, n = 3 mice per genotype, one-way ANOVA, Tukey's multiple comparisons.

(C) Percentage of silent (0 $Ca^{2+}$-transients per min) neurons in motor cortex layer 5 of WT, (GR)400, and (PR)400 mice at 15-19 months of age. Graph, mean ± SEM, n = 3 mice per genotype, one-way ANOVA, Tukey's multiple comparisons.

(D) Mean firing rate ($log_{10}$Hz) of neurons motor cortex layer 5 in WT, (GR)400, and (PR)400 mice at 15-19 months of age; black lines indicate medians, dashed lines indicate quartiles. Graph, n = 3 mice per genotype, one-way ANOVA, Tukey's multiple comparison.

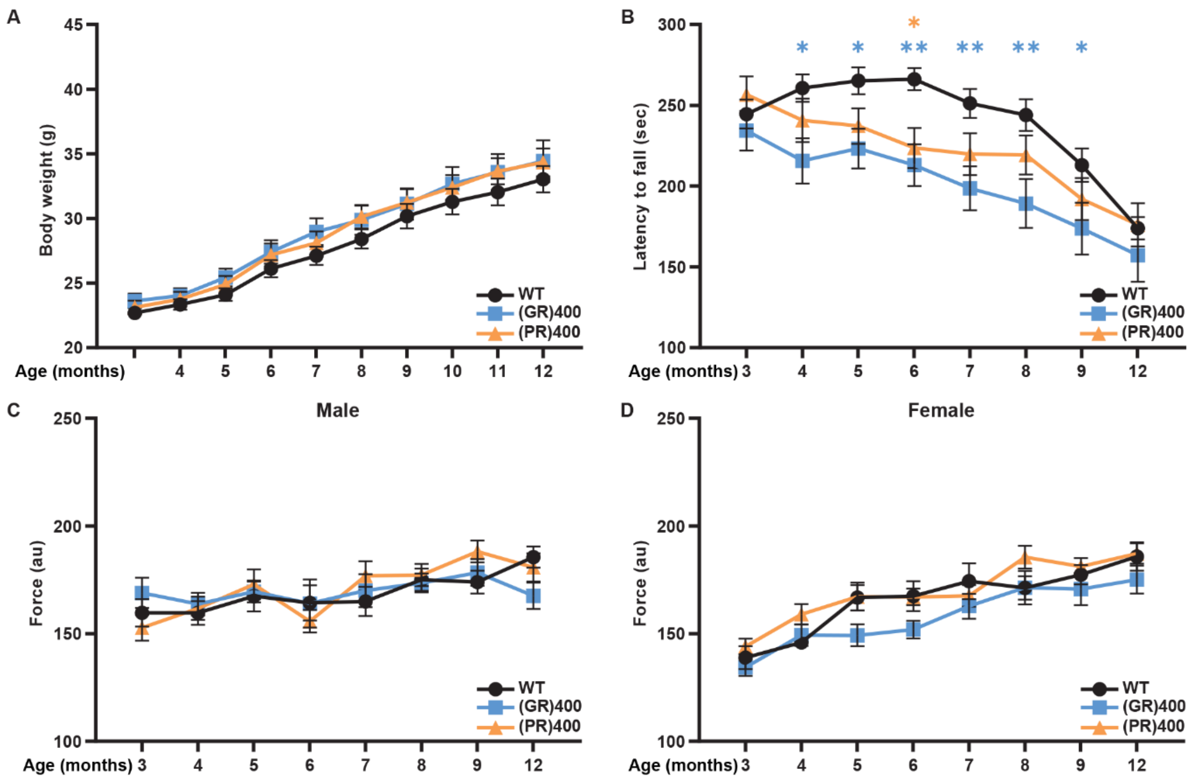


Supplementary Figure 5. **Female poly(GR) and poly(PR) knock-in mice develop rotarod impairment**

(A) Body weights of WT, (GR)400, and (PR)400 mice up to 12 months of age. Graph, mean ± SEM, n = 14 mice per genotype, two-way ANOVA, Bonferroni's multiple comparison.

(B) Accelerated rotarod analysis of motor coordination in WT, (GR)400, and (PR)400 mice up to 12 months of age. Graph, mean ± SEM, n = 14 mice per genotype, two-way ANOVA, Bonferroni's multiple comparison, $*p < 0.05$, $**p < 0.01$.

(C) Grip strength analysis of muscle force in WT, (GR)400, and (PR)400 male mice over lifespan. Graph, mean ± SEM, n = 14 mice per genotype, two-way ANOVA, Bonferroni's multiple comparison.

(D) Grip strength analysis of muscle force in WT, (GR)400, and (PR)400 female mice over lifespan. Graph, mean ± SEM, n = 14 mice per genotype, two-way ANOVA, Bonferroni's multiple comparison.

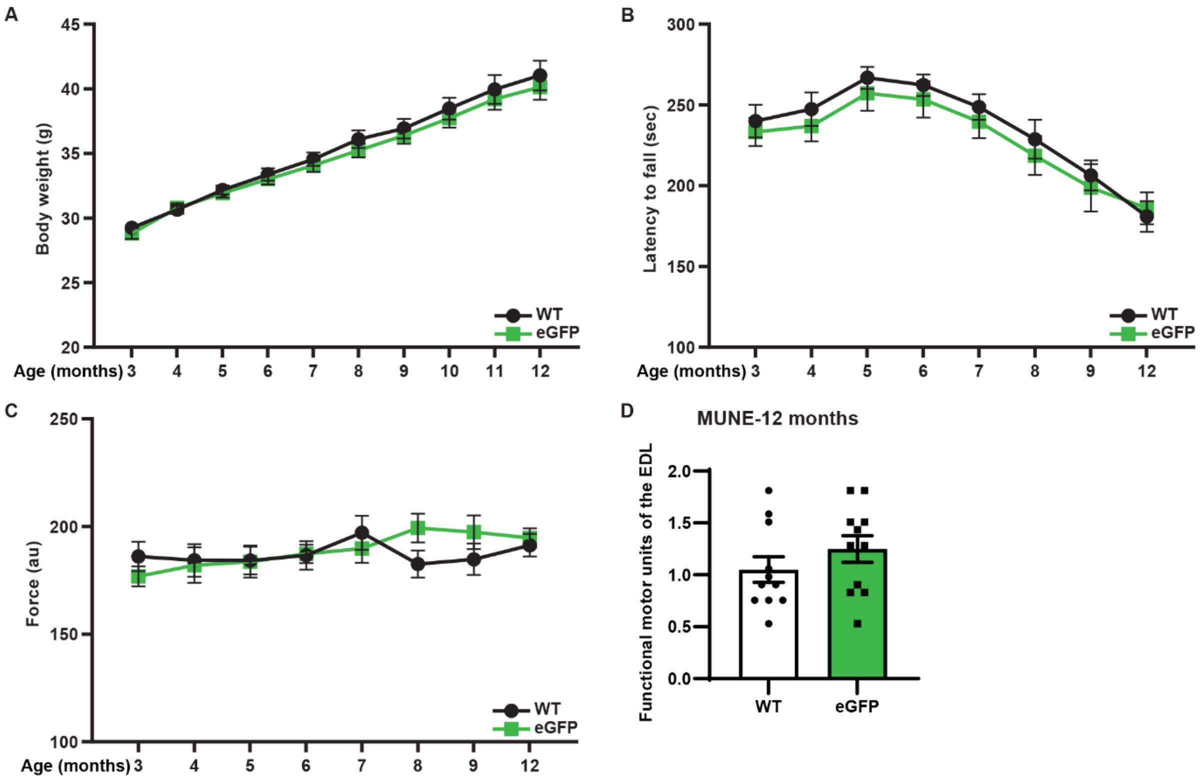


Supplementary Figure 6. **eGFP knock-in mice do not develop motor unit reduction or rotarod impairment**

(A) Body weight analysis of WT and eGFP mice over lifespan. Graph, mean ± SEM, n = 14 mice per genotype, two-way ANOVA, Bonferroni's multiple comparison.

(B) Accelerated rotarod analysis of motor coordination in WT and eGFP mice over lifespan. Graph, mean ± SEM, n = 14 mice per genotype, two-way ANOVA, Bonferroni's multiple comparison.

(C) Grip strength analysis of muscle force in WT and eGFP mice over lifespan. Graph, mean ± SEM, n = 14 mice per genotype, two-way ANOVA, Bonferroni's multiple comparison.

(D) Quantification of MUNE determined in EDL muscle in WT and eGFP mice at 12 months of age. Graph, mean ± SEM, n mice = 6 WT and 6 eGFP, n muscles = 11 WT and 11 eGFP, unpaired two-sample Student's t-test.

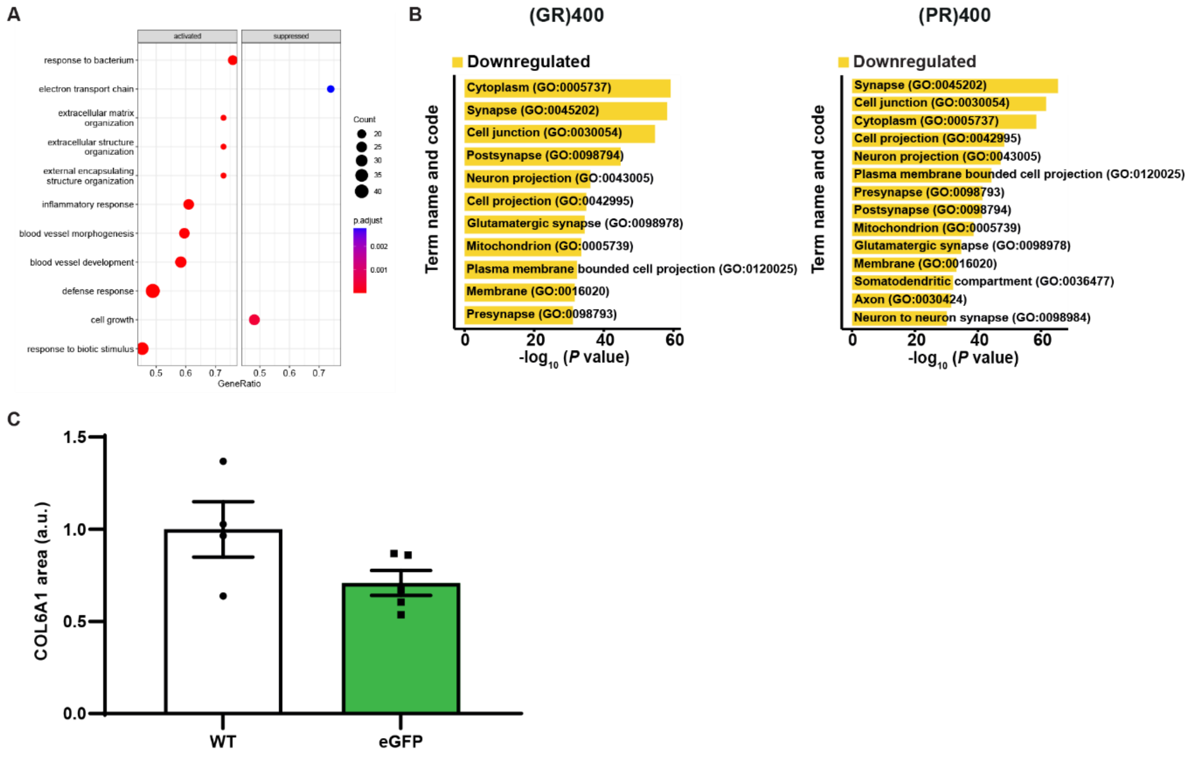


Supplementary Figure 7. **(GR)400 and (PR)400 knock-in mouse spinal cord extracellular matrix signature is conserved in *C9orf72* patient motor neurons and (GR)400 and (PR)400 knock-in mouse spinal cord has decreased synapse protein levels**

(A) Significantly enriched activated (left) and suppressed (right) Gene Ontology (GO) pathways in *C9orf72* patient laser capture microdissected spinal cord motor neuron microarray data. The names of GO terms are shown on the x axis, while the y axis represents the gene ratio. The depth of the color illustrates the adjusted p-value. The size of the circle in the graph corresponds to the size of the gene set.

(B) GO analysis displaying terms name and code from significantly downregulated (yellow) proteins in the lumbar spinal cord of (GR)400 (left panel), or (PR)400 (right panel) mice at 12 months of age. Proteomics performed on n = 5 mice per genotype.

(C) Quantification of immunofluorescence staining of COL6A1 in lumbar spinal cord in WT and eGFP mice at 12 months of age. Graph, mean ± SEM, n = 4 mice per genotype, unpaired two-sample Student's t-test.
